# Pericyte-derived fibrotic scarring is conserved across diverse central nervous system lesions

**DOI:** 10.1101/2020.04.30.068965

**Authors:** David O. Dias, Jannis Kalkitsas, Yildiz Kelahmetoglu, Cynthia P. Estrada, Jemal Tatarishvili, Aurélie Ernst, Hagen B. Huttner, Zaal Kokaia, Olle Lindvall, Lou Brundin, Jonas Frisén, Christian Göritz

## Abstract

Fibrotic scar tissue limits central nervous system regeneration in adult mammals. The extent of fibrotic tissue generation and distribution of stromal cells across different lesions in the brain and spinal cord has not been systematically investigated in mice and humans. Furthermore, it is unknown whether scar-forming stromal cells have the same origin throughout the central nervous system and in different types of lesions. In the current study, we compared fibrotic scarring in human pathological tissue and corresponding mouse models of penetrating and non-penetrating spinal cord injury, traumatic brain injury, ischemic stroke, multiple sclerosis and glioblastoma. We show that the extent and distribution of stromal cells are specific to the type of lesion and, in most cases, similar between mice and humans. Employing *in vivo* lineage tracing, we report that in all mouse models developing fibrotic tissue, the primary source of scar-forming fibroblasts is a discrete subset of perivascular cells, termed type A pericytes.

We uncover pericyte-derived fibrosis as a conserved mechanism that may be explored as a therapeutic target to improve recovery after central nervous system lesions.

## Introduction

Regeneration of the central nervous system (CNS) in adult mammals is limited. Scar tissue that forms after CNS lesions is essential for sealing off the injured tissue and containing the damage^1–4^. However, it also functions as a permanent barrier to regenerating axons, contributing to the long-lasting functional deficits observed after injury^5,6^.

The mature CNS scar is a compartmentalized, multicellular structure. Non-neural cells, including extracellular matrix (ECM)-producing stromal fibroblasts and peripherally-derived macrophages, constitute the fibrotic, or stromal, component of the scar^2^. Most research has focused on reactive astrocytes and NG2-expressing glia constituting the glial component of the scar^5,7^, and much less is known about the fibrotic scar tissue.

Using *in vivo* lineage tracing, we previously identified a small subset of perivascular cells, named type A pericytes, as the cellular origin of stromal fibroblasts present in fibrotic scar tissue after penetrating spinal cord injury. Type A pericytes are initially recruited to mediate wound closure, but consequently form fibrotic scar tissue, which constitutes a barrier for regeneration^6^. Type A pericytes are located in the vascular wall encased by basal lamina and represent about 10% of all platelet-derived growth factor receptor-beta (PDGFRβ)-expressing pericytes in the uninjured adult mouse spinal cord. They can be distinguished from other non-scar forming pericytes by the expression of the glutamate aspartate transporter GLAST (EAAT1), which can be used for selective targeting of type A pericytes. Following penetrating spinal cord injuries, type A pericytes enter the lesion site with sprouting vessels during a transient phase of vascular remodeling. Between 3 and 9 days post-injury, type A pericytes proliferate intensely, break through the surrounding vascular basal lamina and migrate into the lesion, where they cluster and contribute to the formation of the fibrotic scar^3^. Although different cell types were proposed to contribute^8–11^, the cellular origin of scar-forming fibroblasts following CNS lesions in the brain remains elusive, due to the lack of *in vivo* genetic fate mapping studies required to unequivocally trace cells and establish lineage relationships. It is also unclear to what extent fibrotic scar tissue enriched in stromal fibroblasts is formed in humans after CNS lesions, such as spinal cord injury, stroke, multiples sclerosis (MS) and glioblastoma multiforme (GBM).

In the present study, we performed *in vivo* lineage tracing and explored the contribution of type A pericytes to fibrotic scarring in mouse models of penetrating and non-penetrating spinal cord injury, traumatic brain injury, ischemic stroke, MS and GBM. We report that fibrotic scar tissue formation by pericytes is preserved throughout the CNS, but the extent and distribution of stromal cells varies depending on the lesion. Type A pericytes are the main source of PDGFRβ-expressing stromal cells in mouse models of penetrating and non-penetrating spinal cord injuries, traumatic brain injury, ischemic stroke and MS, but contribute less extensively to tumor stroma.

Furthermore, we found that humans develop non-neural, fibrotic scar tissue with large amounts of non-vessel associated stromal fibroblasts after spinal cord injury. In spinal cords of individuals with active MS, PDGFRβ–expressing stromal cells mostly accumulated in the perivascular space. In comparison, PDGFRβ–expressing stromal cells increased in density but remained mostly associated with the vasculature in the ischemic lesion core of stroke patients and in the stroma of aggressive grade IV human GBM tumors.

## Results

### Genetic labeling of type A pericytes in the adult mouse CNS

We employed GLAST-CreER^T2^ transgenic mice^12^ carrying a Rosa26-enhanced yellow fluorescent protein (R26R-EYFP) reporter allele (hereafter referred to as GLAST-CreER^T2^;R26R-EYFP) to genetically fate map type A pericytes, as previously established in the spinal cord^3,6^ (Supplementary Fig.1a). Upon tamoxifen-mediated genetic recombination, GLAST-expressing type A pericytes are inheritably labelled by EYFP expression. Under homeostatic conditions, type A pericytes are associated with blood vessels throughout the grey and white matter spinal cord parenchyma and express the accepted pericyte markers PDGFRβ and CD13 (also known as aminopeptidase N) (Supplementary Fig. 1b-d), but are not marked by desmin and alpha smooth muscle actin (αSMA), found in other mural cells (*i.e*., type B pericytes and vascular smooth muscle cells)^3^. Likewise, we detected EYFP-expressing perivascular cells sharing the same marker expression and comprising about 10% of all PDGFRβ-expressing cells in the uninjured cerebral cortex and striatum (Supplementary Fig. 2a-c), identifying them as type A pericytes. Apart from type A pericytes, we observed occasional recombination in a small subset of ependymal cells and white matter radial astrocytes in the spinal cord^3^, and associated with the vasculature in the meninges surrounding the brain and spinal cord (Supplementary Fig. 1b and 2a). Additionally, the GLAST-CreER^T2^ transgene allows recombination in a large subset of parenchymal astrocytes throughout the brain (*i.e*., striatal and cortical astrocytes, Supplementary Fig. 2d), as well as in subventricular zone astrocyte-like neural stem cells^12,13^. Astrocytes, ependymal cells and their respective progeny do not express the stromal marker PDGFRβ, which allows distinguishing them from pericyte-derived scar-forming fibroblasts.

### Type A pericytes contribute to fibrotic scar formation following penetrating and non-penetrating spinal cord injury

To compare the contribution of type A pericytes to fibrotic scarring after penetrating and non-penetrating spinal cord injury, we lineage traced type A pericytes after a dorsal *funiculus* incision^3,14,15^ and after a complete crush injury^16^, respectively (Fig. 1 and Supplementary Fig. 3). Injuries were performed following a clearing period of 7 days after tamoxifen-mediated recombination, ensuring that all recombination occurs before lesioning^3,6^.

**Figure 1.**
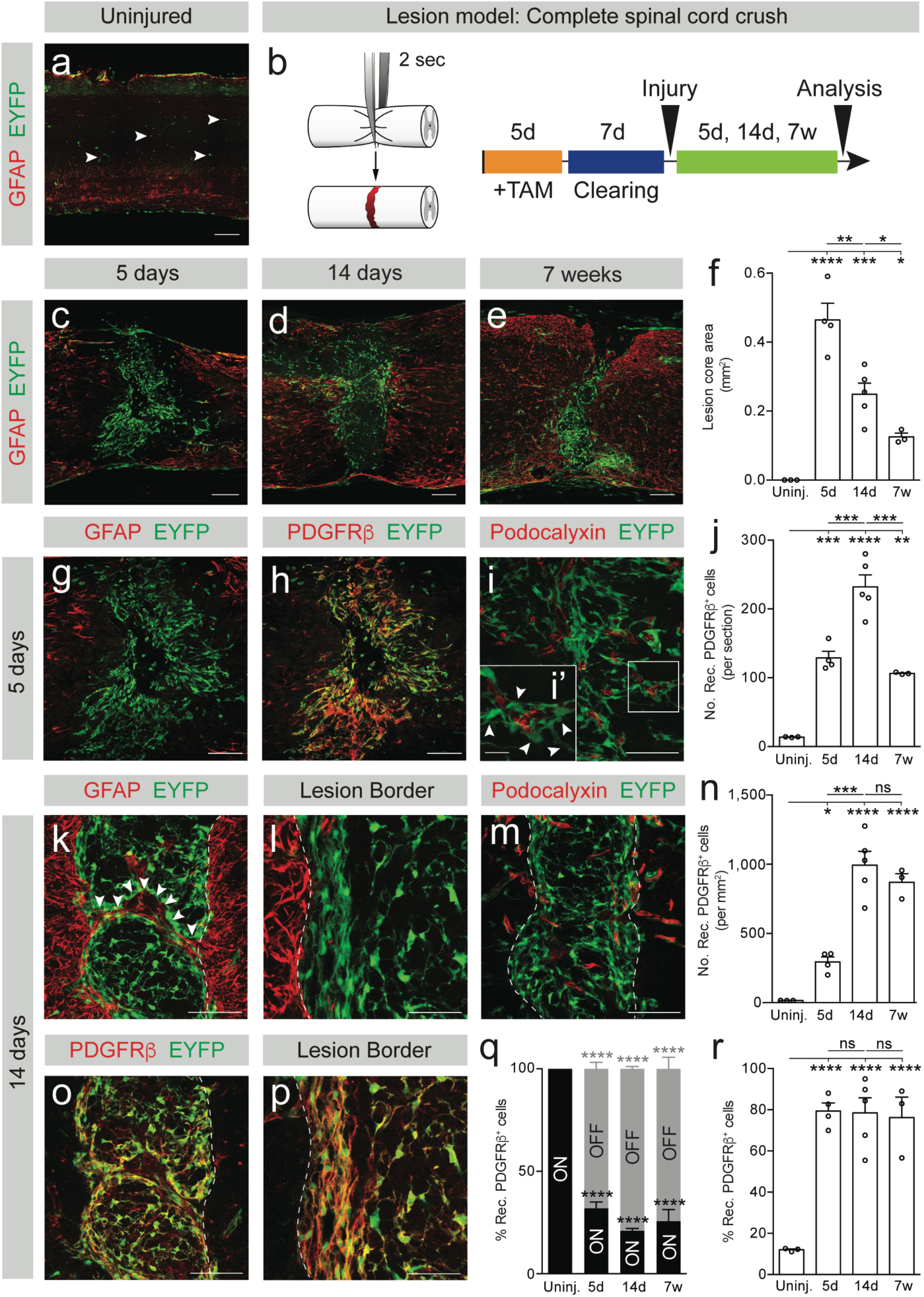
Type A pericyte progeny is the main source of stromal fibroblasts that form fibrotic scar tissue after non-penetrating spinal crush injury. **(a)** In uninjured GLAST-CreER^T2^; R26R-EYFP mice, a subset of perivascular cells lining blood vessels, named type A pericytes, are recombined (EYFP^+^, arrowheads) and dispersed throughout the spinal cord grey and white matter. **(b)** Spinal cord complete crush injury model and experimental timeline. (**c**-**e**) Distribution of recombined cells at 5 days (c), 14 days (d) and 7 weeks (e) following spinal cord injury. (**f**) The lesion core shrinks over time as the scar matures. (**g**-**i**) Recombined cells, co-expressing the stromal marker PDGFRβ, enter the lesion area demarcated by GFAP^+^ reactive astrocytes (g, h) and are located in distance to the blood vessel wall (endothelial cells marked by podocalyxin) at 5 days post-injury (i). Arrowheads in (i’) point at recombined cells that are located outside the vascular wall. (**j**) The number of recombined PDGFRβ^+^ cells peaks at 14 days post-injury and decreases at more chronic stages. (**k-m**) A sharp border segregating the glial (GFAP^+^) and fibrotic (type A pericyte-derived, EYFP^+^) compartments of the scar forms at 14 days post-injury (k,l). A fraction of EYFP^+^ cells is located away from the blood vessel wall (m). Arrowheads in k point at a glial bridge traversing the lesion core. (**n**) The density of recombined PDGFRβ^+^ cells populating the fibrotic core increases overtime and remains constant from 14 days after injury onwards. (**o**,**p**) Recombined cells express the stromal marker PDGFRβ and populate the core of the lesion 14 days post-injury (o). EYFP^+^ cells taking part in the fibrotic-glial border (**l**) orient differently than EYFP^+^ cells found in the inner core of the scar (p). **(q)** Under homeostatic conditions, all recombined PDGFRβ^+^ cells are found associated with the blood vessel wall (ON vessel). After injury, a fraction recombined is located away from the vascular wall (OFF vessel). **(r)** Under homeostatic conditions, type A pericytes comprise about 10% of the total PDGFRβ-expressing pericytes. After injury, type A pericyte-derived cells are the main contributors to PDGFRβ^+^ stromal cells populating the lesion core. TAM, tamoxifen; GFAP, glial fibrillary acidic protein; EYFP, enhanced yellow fluorescent protein; PDGFRβ, platelet-derived growth factor-beta. Scale bars represent 200 µm (a,c-e), 100 µm (g-i, k,m,o), 50 µm (l, p) and 25 µm (i’). Data shown as mean ± s.e.m. n=3 (Uninjured), n=4 (5 dpi), n=5 (14 dpi) and n=3 (7 wpi). ns, non-significant; \**P*<0.05, **P<0.01, ***P<0.001, \*\*\*\**P*<0.0001 by One-Way ANOVA followed by Holm-Sidak *post hoc* test. g, h, and k,o, and l,p denote paired images. All images show sagittal sections.

We compared type A pericyte-derived scar formation during the sub-acute phase at 5 days post-injury (dpi), at the time of glial-stromal border establishment at 14 dpi, and at a chronic time point at 6-7 weeks post-injury (wpi) (Fig. 1a-f and Supplementary Fig. 3a-e). While the uninjured spinal cord only contained a small number of recombined (EYFP^+^) type A pericytes (Fig. 1a, Supplementary Fig. 1 and Supplementary Fig. 3b), we found many EYFP^+^ type A pericyte-derived cells within the lesioned parenchyma at all studied time points after complete spinal crush and dorsal *funiculus* incision (Fig. 1c-e and Supplementary Fig. 3c-e). At 5 dpi, the total number of EYFP^+^ cells co-expressing the stromal marker PDGFRβ had drastically increased in the lesion core compared to uninjured conditions, and most recombined cells were no longer associated with the blood vessel wall (Fig. 1g-j and Supplementary Fig. 3f-h). The number of EYFP^+^PDGFRβ^+^ cells peaked at 14 dpi and then dropped by 6-7 wpi, as the scar matured (Fig. 1j and Supplementary Fig. 3l). Over time, the lesion site condensed (Fig. 1f and Supplementary Fig. 3i), leading to increased EYFP^+^ cell density in the lesion core (Fig. 1n and Supplementary Fig. 3m). At 14 dpi the fibrotic lesion core was flanked by GFAP^+^ astrocytes, establishing a sharp border between the glial and fibrotic scar compartments (Fig. 1k,l). We noticed that type A pericyte-derived cells apposed to reactive glia at the glial-fibrotic lesion border exhibited a distinct morphology and arranged differently compared to EYFP^+^ cells in the inner core of the fibrotic scar (Fig. 1l,p). In scar regions devoid of pericyte-derived cells, we could observe GFAP^+^ astrocyte processes crossing the lesion site, forming glial bridges (Fig. 1k,o).

Fibrotic scar tissue contains PDGFRβ^+^ stromal cells associated with the blood vessel wall (ON the vessel), representing perivascular cells, and extravascular (OFF the vessel) PDGFRβ^+^ stromal cells, representing fibroblasts. We noticed that spinal cord injury triggered robust detachment of recombined stromal cells from the blood vessel wall, as approximately 56±4% (dorsal *funiculus* injury) and 74±6% (crush spinal cord injury) of EYFP^+^/PDGFRβ^+^ cells were no longer associated with the blood vessel wall at chronic stages (Fig.1q and Supplementary Fig.3n). After complete spinal cord crush injury, the overall contribution of type A pericytes to all PDGFRβ^+^ stromal cells was on average 80% at all time points investigated. This value is likely to represent an underestimate as incomplete recombination is expected^3,6^ and the highest animal showed up to 95% contribution (Fig. 1r).

Type A pericytes reacted in a similar manner and showed comparable dynamics in response to penetrating and non-penetrating lesions to the spinal cord (Fig.1 and Supplementary Fig. 3). We did not observe significant contribution from sparsely recombined ependymal cells and radial astrocytes to scar-forming cells.

We conclude that nearly all scar-forming PDGFRβ^+^ stromal cells derive from type A pericytes after penetrating and non-penetrating injuries to the spinal cord.

### Type A pericytes contribute to fibrotic scar formation following traumatic brain injury

To investigate whether type A pericyte-derived scarring is conserved throughout the CNS, we lineage traced type A pericytes after a stab wound to the brain (Fig. 2 and Supplementary Fig. 4a).

**Figure 2.**
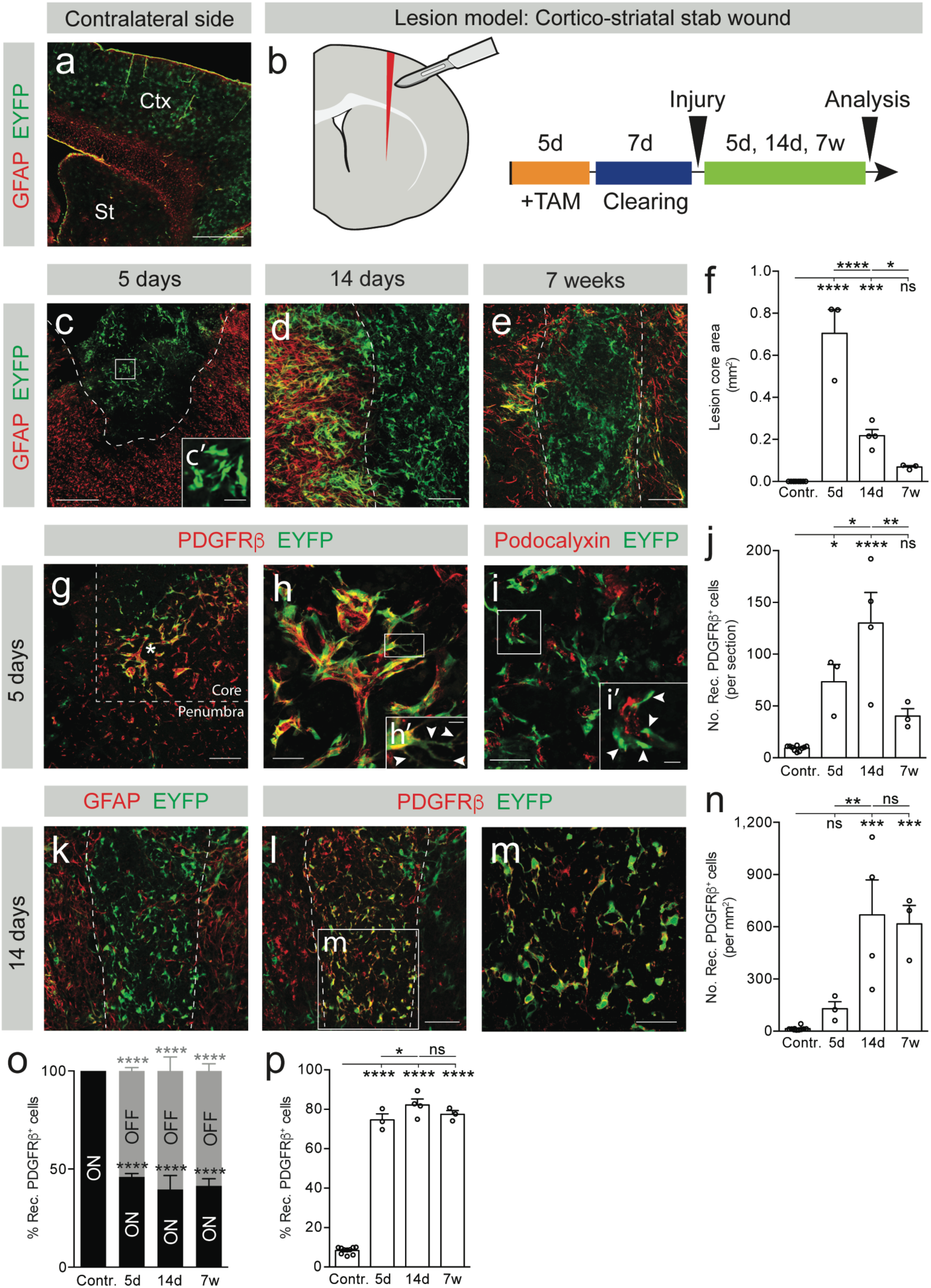
Type A pericyte progeny is the main source of stromal fibroblasts that form fibrotic scar tissue after cortico-striatal stab lesions. **(a)** Type A pericytes and parenchymal astrocytes are recombined (EYFP^+^) throughout the cerebral cortex and striatum of GLAST-CreER^T2^; R26R-EYFP mice. **(b)** Cortico-striatal stab lesion model and experimental timeline. (**c**-**e**) Distribution of recombined cells at 5 days (c), 14 days (d) and 7 weeks (e) following injury. (c’) shows recombined cells with fibroblast-like morphology present in the lesion core. (**f**) The lesion core area reduces over time, as the scar matures. (**g**-**i**) PDGFRβ-expressing recombined cells concentrate towards the core of the lesion (g, h) and a fraction is located outside the vascular wall (endothelial cells marked by podocalyxin) at 5 days post-injury (i). Arrowheads in (h’) and (i’) point at protrusion-like structures and recombined cells that had lost contact with the vascular wall, respectively. Asterisk in (g) marks the lesion core. (**j**) The number of recombined PDGFRβ^+^ cells peaks by 14 days after injury and declines thereafter, as the scar matures. (**k-m**) Recombined cells, co-expressing the stromal marker PDGFRβ, occupy the lesion core and are bordered by reactive glia at 14 days post-injury. **(n)** The density of recombined PDGFRβ^+^ cells at the fibrotic lesion core increases with time and remains constant from 14 days post-injury onwards. **(o)** All recombined cells are found attached to the vascular wall (ON vessel) in the contralateral side to the lesion. After injury, a fraction of recombined PDGFRβ^+^ cells is located outside the vessel wall (OFF vessel). **(p)** Type A pericytes comprise about 10% of all PDGFRβ-expressing pericytes in the cerebral cortex and striatum in the contralateral side to the lesion. Following injury, type A pericyte-derived cells are the main contributors to PDGFRβ^+^ stromal cells found at the lesion core. Ctx, cortex; St, striatum. Scale bars represent 500 µm (a), 400 µm (c), 200 µm (g), 100 µm (d,e,k,l), 50 µm (c’, h, i, m) and 10 µm (h’, i’). Data shown as mean ± s.e.m. n=9 (Contralateral), n=3 (5 dpi), n=3 (14 dpi) and n=3 (7 wpi). ns, non-significant; \**P*<0.05, **P<0.01, ***P<0.001, \*\*\*\**P*<0.0001 by One-Way ANOVA followed by Holm-Sidak *post hoc* test. (k, l) denote paired images. All images show sagittal sections.

Discrete stab lesions restricted to the cerebral cortex mainly induce gliosis^17^ and do not generate extensive fibrotic tissue, as seen by no substantial increase in PDGFRβ^+^ stromal fibroblasts at 14 dpi (Supplementary Fig. 4b-f). As expected, we observed recombined GFAP^+^ astrocytes (Supplementary Fig. 4b) but little to no contribution of lineage traced type A pericytes after cortical stab lesions (Supplementary Fig. 4c-f). In contrast, larger cortico-striatal stab lesions triggered robust fibrotic and glial scarring (Fig. 2a-e) and allowed to investigate the contribution of type A pericytes to fibrotic scarring in the brain. At 5 dpi, we found type A pericyte-derived EYFP^+^ cells co-expressing the stromal marker PDGFRβ dispersed over the lesion core and no longer associated with the blood vessel wall (Fig. 2c and 2f-i). The number of recombined PDGFRβ^+^ stromal cells peaked at 14 dpi and then dropped by 7 wpi (Fig. 2j). At 14 dpi a compact fibrotic scar core filled with type A pericyte-derived stromal cells was walled off by GFAP-expressing, partially recombined astrocytes (Fig. 2d,k-m) and remained up to 7 wpi (Fig. 2e). Overtime, the scar matured and condensed, leading to an increase in EYFP^+^ cell density in the lesion core (Fig. 2e,f,n). About half of the recombined stromal cells in the lesion core were no longer associated with the blood vessel wall (Fig. 2o). After a cortico-striatal stab wound around 80% of all PDGFRβ^+^ stromal cells associated with the fibrotic scar were recombined at all time points investigated (Fig. 2p). This number may underrepresent the contribution of type A pericytes as recombination is expected to be incomplete. As for spinal cord injury, we conclude that almost all scar-forming PDGFRβ^+^ stromal cells derive from type A pericytes after cortico-striatal stab lesions.

### Type A pericytes contribute to fibrotic scar formation following experimental autoimmune encephalomyelitis

Next, we explored the contribution of type A pericytes to fibrotic scar tissue formation after experimental autoimmune encephalomyelitis (EAE), a widely accepted model of demyelinating diseases, such as multiple sclerosis (MS) (Fig. 3). Because tamoxifen diminishes the clinical severity of EAE^18^, we extended the clearing time after tamoxifen-induced recombination to 50 days, before inducing EAE in adult GLAST-CreER^T2^;R26R-EYFP transgenic mice (Fig. 3a). For EAE induction, animals were immunized with myelin oligodendrocyte glycoprotein (MOG) peptide emulsified in Complete Freund’s Adjuvant (CFA) and received *Pertussis* toxin. Animals receiving CFA and/or *Pertussis* toxin alone but not immunized with the MOG peptide were used as control. Daily evaluation of clinical symptoms and weight confirmed that the animals manifested signs of chronic disease-related neurological deficits, starting 10 days after immunization (Fig. 3b,c).

**Figure 3.**
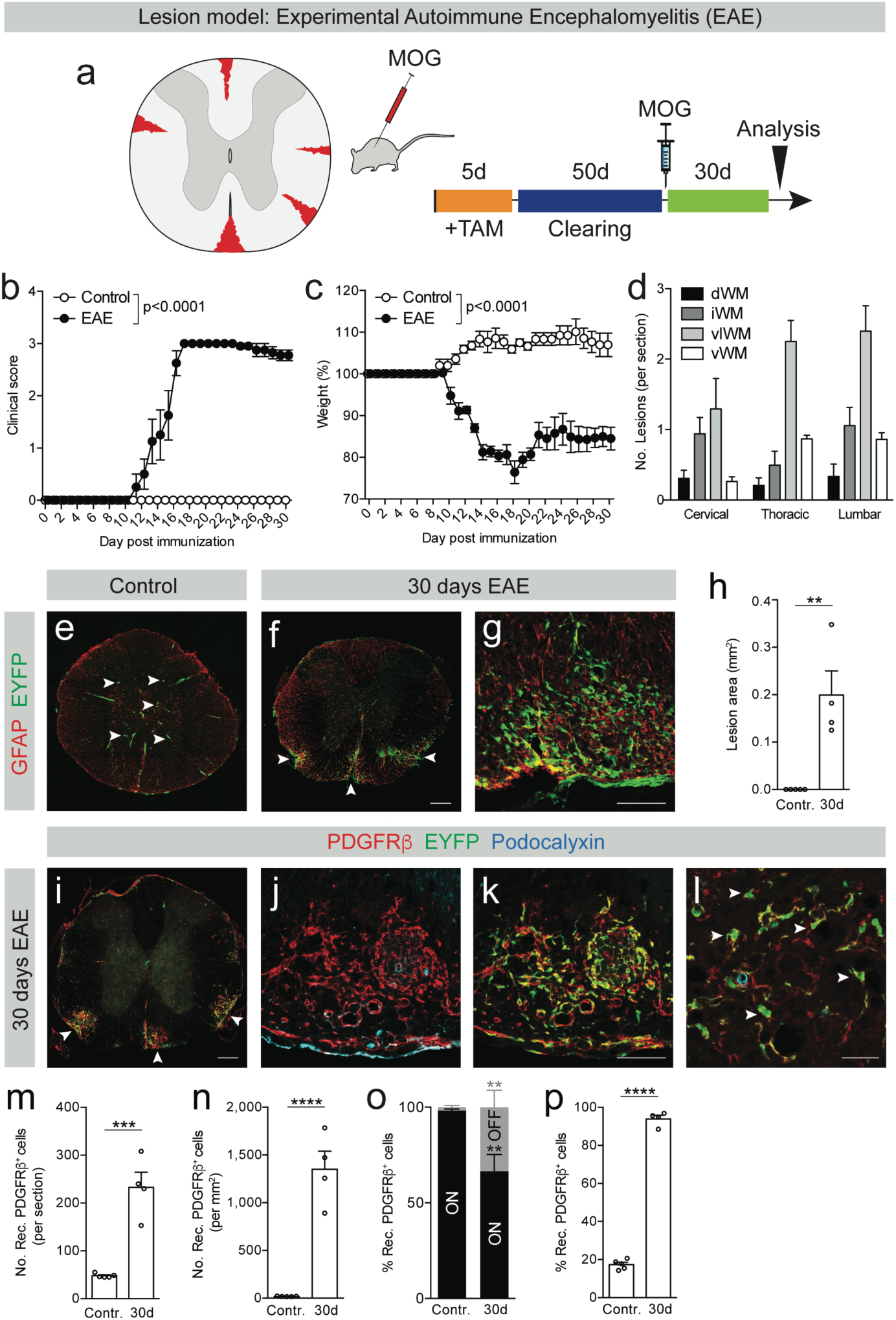
Type A pericyte-derived cells are the main source of fibroblast-like cells making up the fibrotic scar after experimental autoimmune encephalomyelitis. (**a**) MOG_35-55_-induced experimental autoimmune encephalomyelitis model and experimental timeline. **(b**,**c**) Clinical scores and weight curves for EAE and control animals. (**d**) Distribution and number of lesions, 30 days following EAE. (**e, f**) Distribution of recombined cells (arrowheads) in a thoracic segment of a control animal (e) and 30 days following EAE induction (f). **(g)** Reactive astrocytes (GFAP^+^) intermingle with type A pericyte-derived cells in EAE lesions. **(h)** Quantification of the area occupied by EAE lesions. (**i**-**l**) Recombined cells co-expressing the stromal marker PDGFRβ concentrate at the lesion core of EAE scars and a fraction is located in distance to the vascular wall (endothelial cells marked by podocalyxin), 30 days post-injury (i-l). Arrowheads in (i, l) point at EAE lesions and examples of recombined cells that locate away from the vascular wall, respectively. (**m**,**n**) Quantification of the number (m) and density (n) of recombined PDGFRβ^+^ cells in control and EAE animals. **(o)** Recombined PDGFRβ^+^ cells associate with the blood vessel wall (ON vessel) in control animals, whereas after EAE a fraction of recombined PDGFRβ^+^ cells is located in distance to the vascular wall (OFF vessel). **(p)** Type A pericyte-derived cells are the main contributors to PDGFRβ-expressing stromal cells found in EAE scars. MOG, myelin oligodendrocyte glycoprotein. Scale bars represent 200 µm (e,f,i), 100 µm (g, j, k), and 50 µm (l). Data shown as mean ± s.e.m. n=5 (Control) and n=4 (30 dpi). Two-way RM ANOVA: main effect of group, F(1, 8) = 1633, p < 0.0001 in (b) and main effect of group, F(1,8) = 87.48, p < 0.0001 in (c); **p < 0.01, ***p < 0.001, ****p < 0.0001 by two-sided, unpaired Student’s t test in (h) and (m-p). (j,k) denote paired images. All images show coronal sections.

We sacrificed all animals at 30 days post EAE induction, during the chronic disease phase. EAE led to widespread demyelination and scar formation throughout the spinal cord, particularly evident in the ventral and ventrolateral white matter of thoracic and lumbar spinal segments (Fig. 3d,f-h). We did not observe weight loss, disease symptoms or myelin lesions in control animals (Fig. 3b,c,e,h). In all demyelinated lesions we found densely packed recombined PDGFRβ^+^ stromal cells associated with blood vessels and at a distance to them (Fig. 3i-o). Type A pericytes-derived cells accounted for more than 90% of all the PDGFRβ^+^ stromal cells found in demyelinated lesions (Fig. 3p).

In contrast to the crush and stab lesions described before, reactive astrocytes did not sharply wall off the entire lesion, but were instead intermingled with recombined stromal cells within the lesion (Fig. 3f,g). We conclude that fibrotic scar tissue is formed after EAE and that nearly all PDGFRβ^+^ stromal cells are derived from type A pericytes.

### Type A pericytes contribute to stromal cells following ischemic lesions to the brain

We continued to explore whether fibrotic scar tissue formation is preserved in other types of CNS lesions, and investigated the contribution of type A pericytes to an ischemic lesion to the brain (Fig. 4). We induced experimental stroke by a 35 min occlusion of the middle cerebral artery (MCAO), a procedure that generated an ischemic lesion with neuronal loss primarily in the striatum, sparing the cerebral cortex^13^ (Fig. 4b).

**Figure 4.**
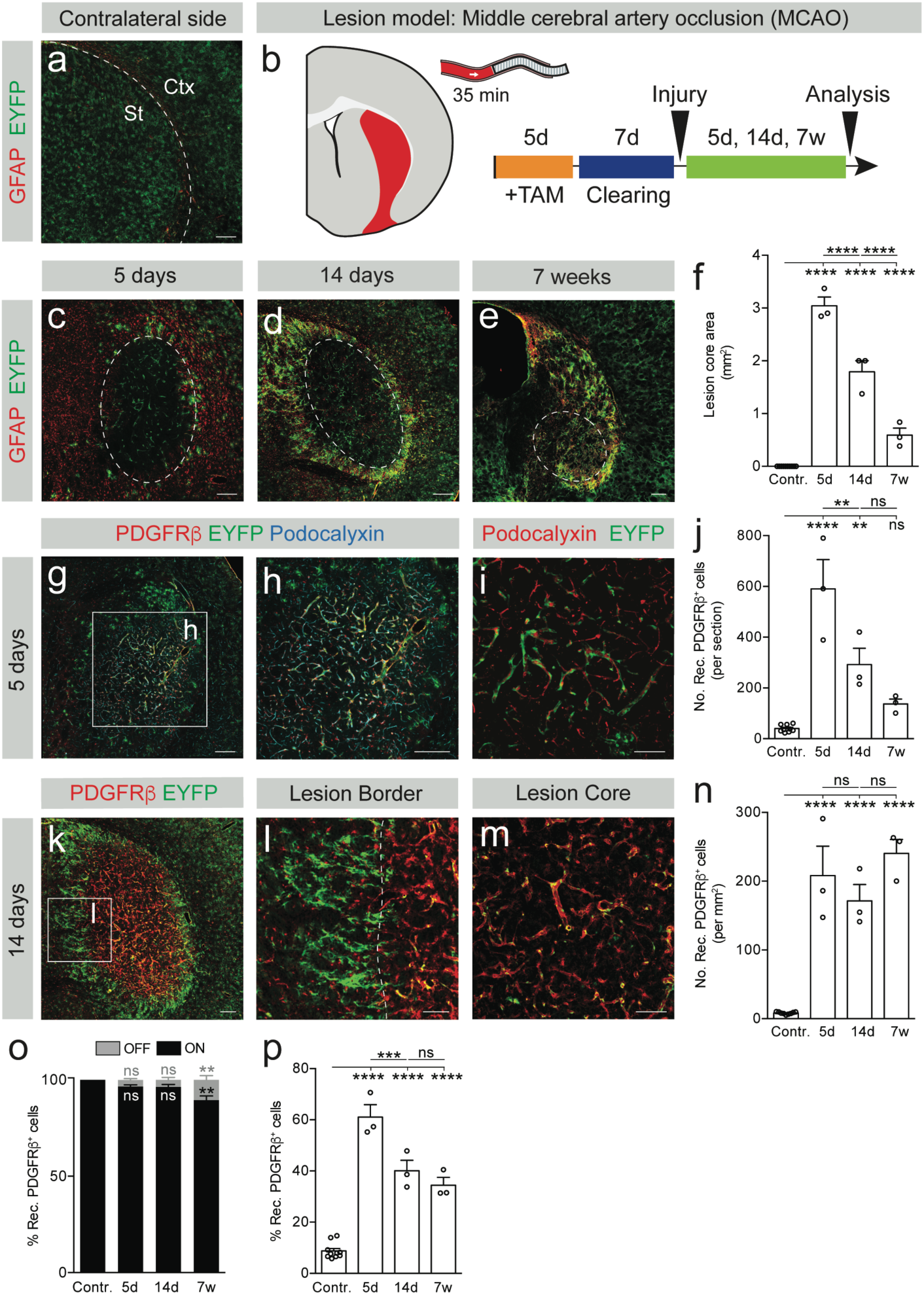
Type A pericytes remain associated with the vascular wall following ischemic stroke confined to the striatum. **(a)** Type A pericytes and parenchymal astrocytes are recombined (EYFP^+^) throughout the cerebral cortex and striatum of GLAST-CreER^T2^;R26R-EYFP mice. **(b)** Middle cerebral artery (MCA) occlusion ischemic stroke model and experimental timeline. Thirty-five minutes occlusion of the MCA results in lesions restricted to the striatum. (**c**-**e**) Distribution of recombined cells at 5 days (c), 14 days (d) and 7 weeks (e) following injury. (**f**) The lesion core contracts over time, as the scar matures. (**g**-**i**) PDGFRβ-expressing recombined cells found in the ischemic stroke core do not leave the vascular wall (endothelial cells marked by podocalyxin) at 5 days post-injury (g-i). (**j**) The number of recombined PDGFRβ^+^ cells peaks by 5 days after injury and regresses subsequently, as the scar matures. (**k-m**) Recombined cells, co-expressing the stromal marker PDGFRβ, occupy the ischemic stroke core at 14 days post-injury (k,l,m). Partially recombined GFAP^+^ reactive glial cells contribute to the glial scar, which surrounds the ischemic core (k, l). **(n)** Injury elicits a sharp increase in the density of recombined PDGFRβ^+^ cells at the ischemic stroke core that remains constant from 5 days to 7 weeks post-injury. **(o)** Most recombined PDGFRβ^+^ cells remain associated with the vascular wall (ON vessel) after striatal ischemic stroke. **(p)** Following injury, recombined PDGFRβ^+^ cells increase in number and represent about half of the total PDGFRβ-expressing stromal cells in the ischemic core. Scale bars represent 200 μm (a,c,d,e,g,k) and 100 μm (h,i,l,m). Data shown as mean ± s.e.m. n=10 (Contralateral), n=3 (5 dpi), n=4 (14 dpi) and n=3 (7 wpi). ns, non-significant; **P<0.01, ***P<0.001, \*\*\*\**P*<0.0001 by One-Way ANOVA followed by Holm-Sidak *post hoc* test. All images show coronal sections.

Type A pericytes were fate mapped as described above for spinal cord injury. At 5 dpi we observed a large number of PDGFRβ^+^ stromal cells in the lesion core. However, in contrast to spinal cord injury and cortico-striatal stab lesions, nearly all PDGFRβ^+^ cells were associated with the vasculature (Fig. 4g,h). Lineage traced type A pericytes and their progeny, co-expressing EYFP and PDGFRβ, had drastically increased in number in the ischemic stroke core at 5 dpi compared to the contralateral stroke side (Fig. 4g-j). Over time, the lesion condensed and the number of recombined cells declined, keeping the cell density stable (Fig. 4 c-f,j,n). At all three time points investigated, nearly all recombined PDGFRβ^+^ cells were associated with the blood vessel wall (Fig. 4g-i, k-m,o). However, while in the contralateral, non-ischemic side, type A pericytes accounted for approximately 10% of all PDGFRβ^+^ cells, 40-60% (depending on the time point) of the PDGFRβ^+^ cells covering blood vessels in the ischemic lesion core were recombined and, therefore, originated from type A pericytes (Fig. 4p). Interestingly, in cases in which the ischemic lesion extended from the striatum into the cortex (cortico-striatal stroke), a large number of PDGFRβ^+^ cells could be observed at a distance from the blood vessel wall, primarily in the cortex (Supplementary Fig. 5). Recombined reactive astrocytes, co-expressing EYFP and GFAP but not PDGFRβ, could be observed in the lesion rim as part of the glial scar (Fig. 4c-e, k,l and Supplementary Fig. 5b).

In summary, we observed that in ischemic lesions confined to the striatum, most of the PDGFRβ^+^ stromal cells are associated with the vascular wall, while after cortico-striatal ischemic stroke a large number of PDGFRβ^+^ stromal cells can also be found in distance to the vessel wall. Type A pericytes contributed substantially to both PDGFRβ^+^ stromal populations.

### Type A pericytes contribute to brain tumor stroma

Next, we explored type A pericyte contribution to tumor stromal tissue formation in the murine glioma 261 (GL261) orthotopic glioma model^19^, which mimics human glioblastoma multiforme (GBM), the most frequent and aggressive malignant brain tumor in adults^20^. For this, we injected mouse GL261 cells in the striatum of adult GLAST-CreER^T2^;R26R-EYFP mice after recombination and tamoxifen clearing (Fig. 5a). Animals developed visible tumors with abnormal and enlarged vasculature, 3 weeks after inoculation (Fig. 5b). The tumor mass covered most of the striatal area and was surrounded by GFAP^+^ astrocytes that were partially recombined (Fig. 5c-e). Compared to the contralateral side to the tumor, the total number and density of type A pericytes in the tumor stroma was reduced (Fig. 5f,g), in line with an overall reduction of pericyte coverage in tumor-associated vessels^21,22^. Nonetheless, we observed that about 30±5% of all type A pericyte-derived EYFP^+^ cells were outside the blood vessel wall and dispersed among tumor cells (Fig. 5h-k).

**Figure 5.**
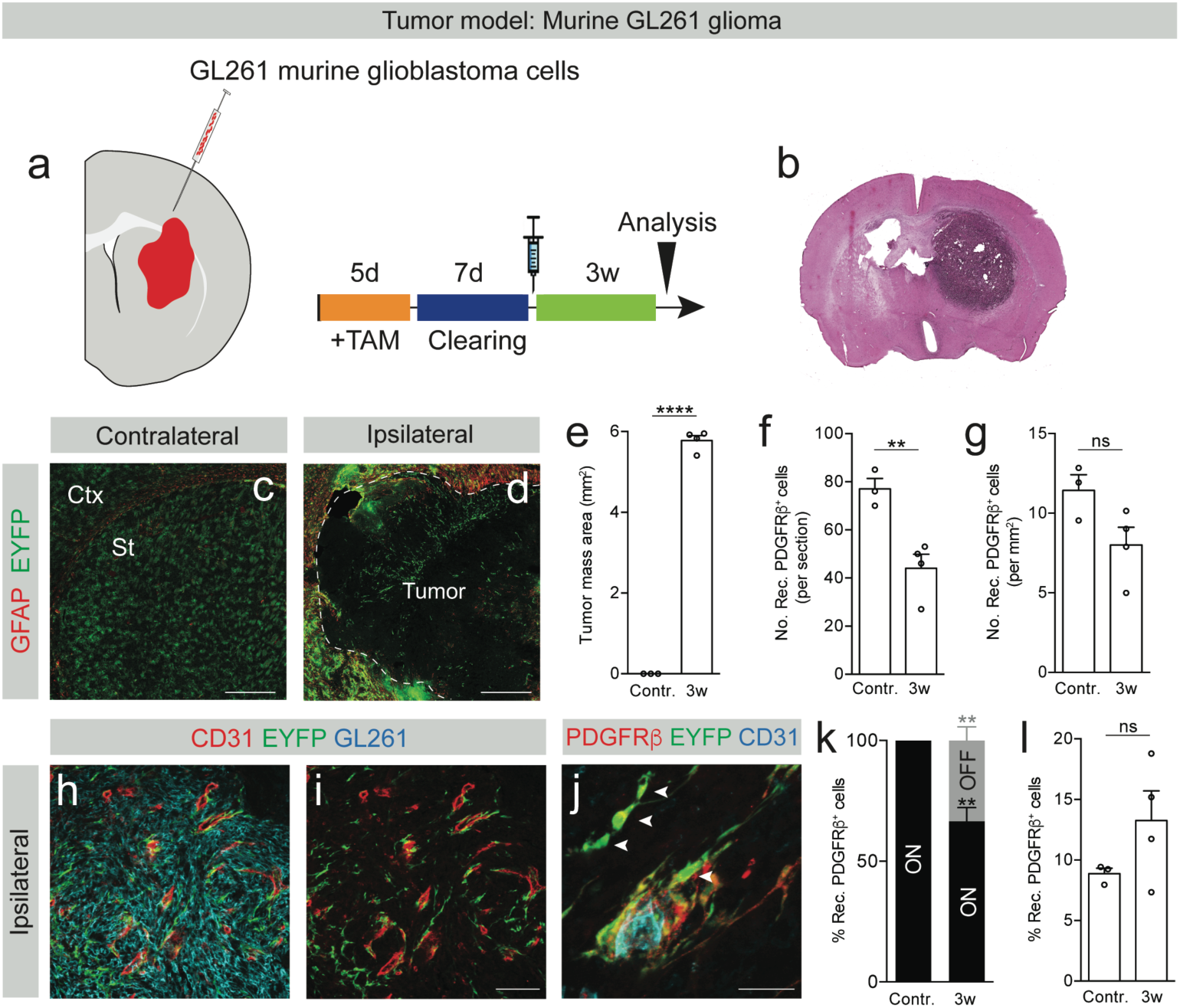
Type A pericyte-derived cells contribute to the tumor stroma. **(a)** Glioblastoma model and experimental timeline. **(b)** Hematoxylin and eosin stain revealed that the tumor cell mass is confined to the striatum. (**c, d**) Distribution of recombined cells in the contralateral (c) and ipsilateral (d) side to the tumor. Recombined pericytes and progeny (EYFP^+^/GFAP^−^) are found within the tumor core (GFAP negative area), whereas partially recombined GFAP^+^ reactive astrocytes contribute to the glial scar bordering the tumor. (**e**) Quantification of the area occupied by the tumor cell mass. (**f, g**) The number (f) and density (g) of recombined PDGFRβ^+^ cells are decreased in the tumor core, compared to the contralateral side of the tumor. (**h**,**i**) Type A pericytes and progeny (EYFP^+^) intermingle with tumor cells. The mouse glioma 261 (GL261) cells are labeled with an antibody against NG2. **(j)** A fraction of recombined cells co-expressing the stromal marker PDGFRβ is found away from the vascular wall (endothelial cells marked by CD31), 3 weeks post-inoculation of tumor cells. Arrowheads point at recombined cells that were located in distance to the blood vessel wall. **(k)** Percentage of recombined PDGFRβ^+^ cells associated (ON vessel) or non-associated (OFF vessel) with the vascular wall in the tumor core. **(l)** Type A pericytes and progeny are not the main source of PDGFRβ-expressing stromal cells found in the core of GL261 murine gliomas. **(j)** Ctx, cortex; St, striatum. Scale bars represent 200 µm (C, D), 100 µm (h,i) and 20 µm (j). Data shown as mean ± s.e.m. n=3 (Contralateral) and n=4 (3 weeks post inoculation). ns, non-significant; **p < 0.01, ****p < 0.0001 by two-sided, unpaired Student’s t-test in (e-g) and (k,l). (h, i) denote paired images. All images show coronal sections.

Overall, type A pericyte-derived cells accounted for 13%±2% of all PDGFRβ^+^ stromal cells in the tumor mass (Fig. 5l). In summary, we found that type A pericytes contribute to tumor-associated PDGFRβ^+^ stromal cells, however not as the main source.

### Fibrotic tissue forms after diverse types of CNS lesions in humans

We investigated the composition of scar tissue that forms following traumatic spinal cord injury in *post-mortem* spinal cord tissue from 6 patients with massive compression or contusion/cyst type injuries in the cervical region, who survived for intervals up to 61 days after injury (Supplementary Table 1). In all cases we found regions of non-neural scar tissue enriched in PDGFRβ^+^ cells, often organized in clusters, and bordered by reactive glia (Fig. 6a,b). The areas of perivascular fibrosis were much larger compared to the surrounding astroglial scar. Significant numbers of PDGFRβ^+^ cells within the lesion core were found at a distance from a blood vessel wall (Fig. 6b inset).

**Figure 6.**
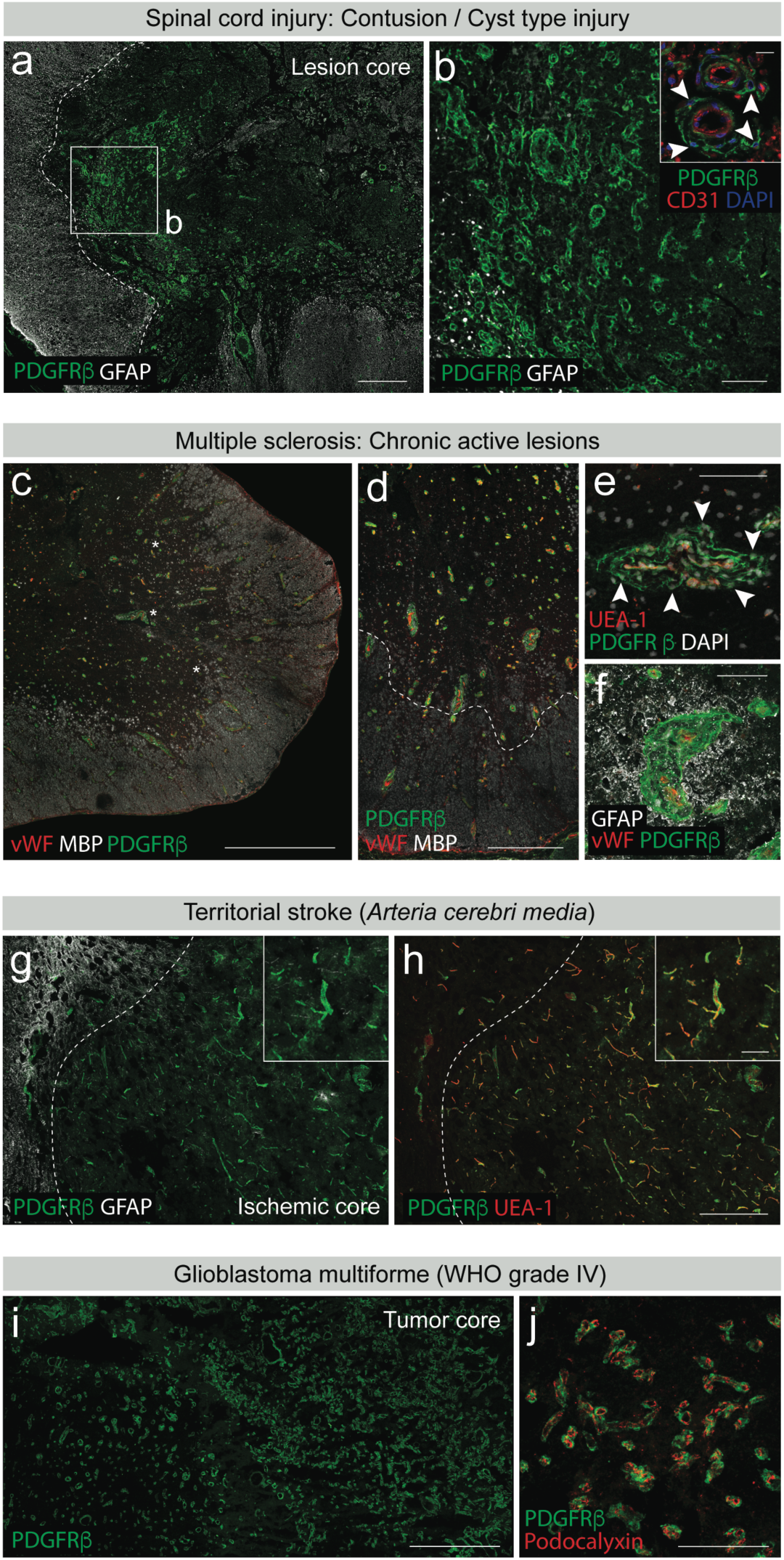
Fibrotic scarring in human pathology. (**a**-**c**) PDGFRβ^+^ stromal cells accumulate at the core of the lesion, 16 days after human traumatic spinal cord injury at C4 (a, b). As observed following experimental spinal cord injury in rodents, a fraction of PDGFRβ^+^ stromal cells is located in distance to the vascular wall (CD31^+^) following traumatic spinal cord injury in humans (inset in b). Dashed line in (a) demarcates the border between GFAP^+^ reactive glia and non-neural scar tissue. (**c**-**f**) Perivascular aggregates of PDGFRβ^+^ stromal cells are detected in chronic active multiple sclerosis lesions (c, d), 21 years following the onset of the disease (secondary progressive multiple sclerosis). Gliosis (GFAP^+^) is observed in regions of PDGFRβ^+^ perivascular cell reactivity (f). Von Willebrand factor marks endothelial cells in c, d and f. Asterisks in (c) denote demyelinated regions (weak MBP signal) of the spinal cord. Arrowheads in E point at PDGFRβ^+^ cells that do not contact the blood vessel wall (UEA-1 marks endothelial cells). Dashed line in (d) separates myelinated white matter tissue (MBP^+^) from partially demyelinated lesion areas. (**g, h**) PDGFRβ^+^ stromal cells remain attached to the blood vessel wall (UEA-1^+^), 7 weeks following territorial stroke in humans (g, h and insets). Dashed line marks the border between GFAP^+^ reactive glia and the ischemic lesion core. (**i, j**) PDGFRβ^+^ stromal cells populate the stroma of a grade IV glioblastoma multiforme that involves the *corpus callosum* and spreads bihemispherically (i). No substantial fraction of PDGFRβ^+^ stromal cells away from the vascular wall (immunopositive for podocalyxin) is observed (j). MBP, myelin basic protein; vWF, von Willebrand factor; UEA-1, *Ulex Europaeus* agglutin I. Nuclei are labeled with 4′,6-diamidino-2-phenylindole (DAPI). Scale bars represent 1000 µm (c, i), 500 µm (a, d, g, h), 200 µm (j), 100 µm (b, e, f, insets in g and h) and 20 µm (inset in b). Error bars show SEM. g, h show paired images. All images show coronal sections.

Likewise, we detected increased perivascular aggregates of PDGFRβ^+^ cells accompanied by gliosis in *post-mortem* spinal cord samples from 6 patients diagnosed with chronic MS (secondary progressive MS type), who survived for 17 to 56 years after the disease onset (Supplementary Table 2, Fig. 6c-f). In active and chronic active MS lesions PDGFRβ^+^ stromal cells were organized in multiple perivascular rings, and a substantial number of cells no longer associated with the blood vessel wall (Fig. 6e). Perivascular fibrosis was closely associated with demyelinated white matter regions. Next, we investigated human *post-mortem* brain tissue obtained from 4 individuals that had a history of a supratentorial territorial or lacunar stroke, involving cerebral cortical and subcortical areas, and who died weeks to months after infarction (Supplementary Table 3). We found that the density of PDGFRβ^+^ cells was elevated in most of the subacute/chronic lesions when compared to surrounding healthy tissue. PDGFRβ^+^ stromal cells within the ischemic lesion core mainly associated with the vasculature (Fig. 6g,h), corroborating our observations following ischemic damage confined to the striatum in the mouse (Fig. 3). The lesion core was flanked by a GFAP^+^ glial scar.

Finally, we investigated brain tissue samples from 3 patients with aggressive grade IV GBM tumors (Supplementary Table 4). All samples presented morphologically abnormal and disorganized vasculature, as opposed to the thin-wall blood vessels of non-malignant adjacent brain tissue. We observed general aberrant hypervascularization within the tumor mass and increased density of PDGFRβ^+^ stromal cells across all 3 patient samples analyzed. Nonetheless, even within the same tumor sample, we found heterogeneity in the density of the blood vessels (Supplementary Table 4 and Fig. 6i). Although malformed blood vessels often presented thick-wall and irregular shape, we found that most of the PDGFRβ^+^ cells within the tumor stroma remained associated with the vasculature (Fig. 6j).

In summary, we observed accumulation of PDGFRβ^+^ stromal cells across all pathological tissues investigated, but their distribution varied depending on the type of lesion. After spinal cord injury and MS, we found accumulation of PDGFRβ^+^ stromal cells in extravascular positions, while we registered an increased number of cells associated with the vasculature after ischemic stroke and in GBM tumors.

## Discussion

We identified a small subset of perivascular cells, termed type A pericytes, as the main source of scar-forming PDGFRβ^+^ stromal cells, contributing to fibrotic tissue generation across a vast number of lesions in the brain and spinal cord. We found that the magnitude of fibrotic scar tissue generation and the distribution of PDGFRβ^+^ stromal cells depend on the extent and type of lesion (Fig. 7). Importantly, the pattern of fibrosis is very similar in mouse models and the corresponding pathology in humans, suggesting that this is an evolutionary conserved mechanism for scar formation in the CNS.

**Figure 7.**
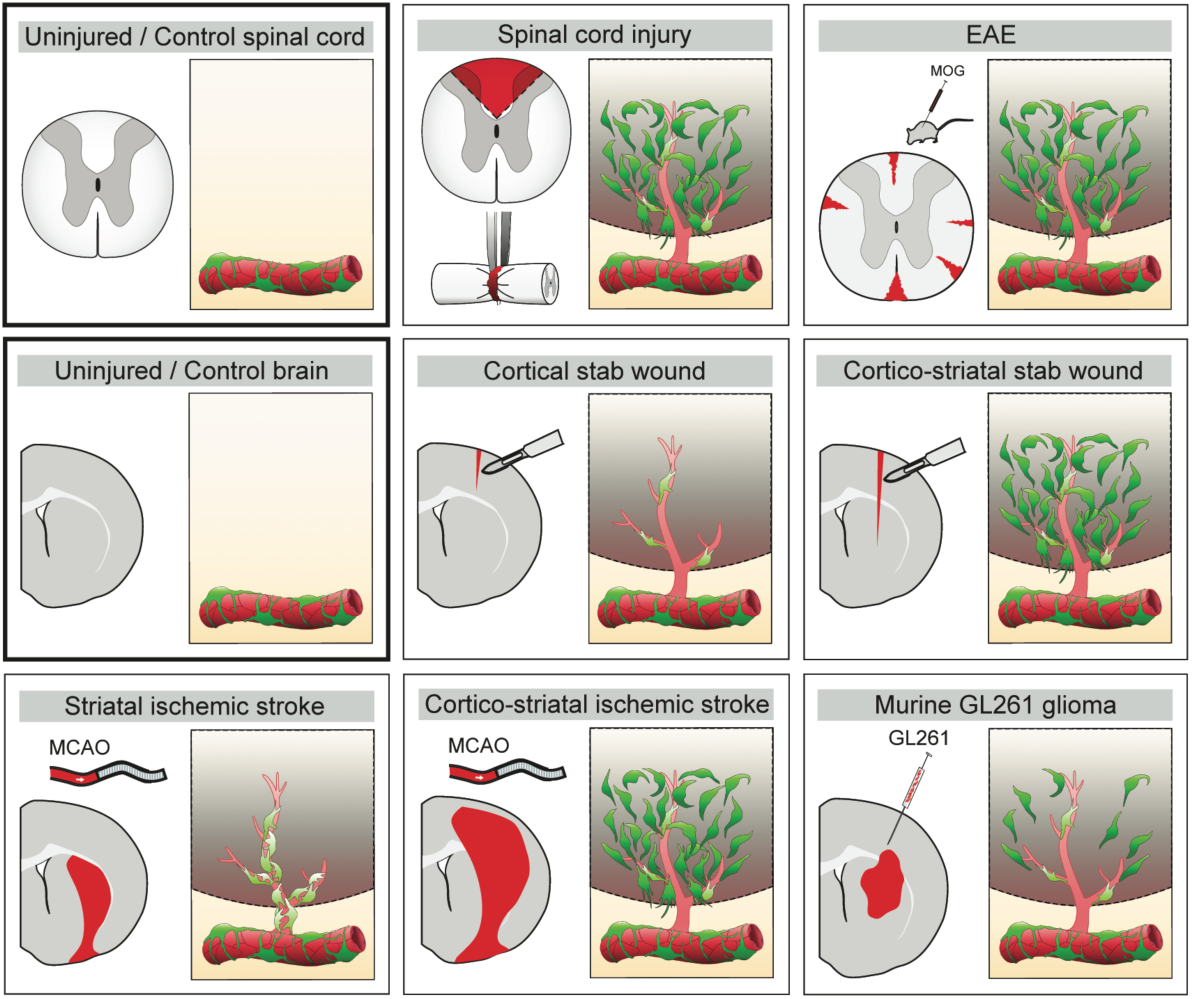
Pericyte-derived fibrotic scar tissue formation is conserved in response to diverse CNS lesions. Schematic illustrations depicting the contribution of type A pericyte-derived cells to diverse CNS lesions. In the uninjured/control brain and spinal cord, all type A pericytes are found associated with the blood vessel wall. After penetrating and non-penetrating spinal cord injury, and EAE, type A pericytes are recruited from the vascular wall and give rise to progeny that form fibrotic scar tissue. Stab lesions restricted to the cerebral cortex do not generate extensive fibrotic scar tissue and, therefore, little to no participation of type A pericytes is observed. However, in larger cortico-striatal stab lesions, type A pericytes give rise to progeny that locate away from the vascular wall and cluster at the core of the lesion, contributing to fibrotic scar tissue formation. Following ischemic stroke confined to the striatum, type A pericytes and progeny increase in number but remain associated with the vascular wall. In contrast, cortico-striatal ischemic lesions trigger type A pericyte recruitment from the blood vessel wall and generation of fibrotic scar tissue by pericyte-derived cells. Although the vasculature of murine GL261 gliomas shows reduced coverage by type A pericytes, some recombined PDGFRβ^+^ stromal cells are found in distance to the blood vessel wall.

Non-penetrating spinal cord injury, in which the *dura mater* remains relatively intact, limits the invasion of *dura*-derived meningeal fibroblasts. Penetrating, and non-penetrating spinal cord injury, led to similar recruitment of PDGFRβ^+^ stromal cells from the vascular wall, and consequent fibrotic scar tissue formation (Fig.1, Fig. 7 and Supplementary Fig. 3). Nearly all PDGFRβ^+^ stromal cells derived from type A pericytes. In humans, we observed extensive fibrotic scar tissue formation enriched in PDGFRβ^+^ stromal cells at subacute stages after massive compression, as well as contusive injuries. These results together with previous reports, acknowledging the accumulation of inflammatory cells and ECM deposits^1,23–26^, show cumulative evidence of fibrotic scar tissue generation after spinal cord injury in humans. In all 6 cases studied, regions of non-neural tissue were much larger compared to the astrocyte scar border (Fig. 6a,b).

Employing a Col1a1-GFP reporter line, Col1a1-expressing perivascular fibroblasts, mainly located at larger vessels, were suggested as the primary source of scar-forming stromal fibroblasts following contusive spinal cord injury^27^ and EAE^28^ in the mouse. Since Col1a1-GFP only detects cells with active Col1a1 transcriptional activity, and no fate mapping of Col1a1-expressing cells has been employed in these studies, the contribution to scar-participating fibroblasts remains speculative.

EAE lesions generated substantial fibrotic scar tissue enriched in PDGFRβ^+^ stromal cells accumulating outside the blood vessel wall (Fig. 3 and Fig. 7). Again, we found that nearly all PDGFRβ^+^ stromal cells derived from type A pericytes. In spinal cords of individuals with active MS lesions, we found PDGFRβ^+^ stromal cells accumulating outside, but in close proximity to the vascular wall (Fig. 6c-f), as reported in MS brain lesions^29^.

In comparison to spinal cord injury and EAE, stab lesions to the mouse brain showed similar fibrotic scar tissue formation only after larger cortico-striatal injuries, but not after smaller injuries, restricted to the cortex (Fig. 2, Fig. 7 and Supplementary Fig. 4). Nearly all PDGFRβ^+^ stromal cells derived from type A pericytes after cortico-striatal stab lesions. A recent study using a Tbx18-CreER^T2^ line to fate map pericytes after discrete cerebral cortex stab lesions led to the generalized conclusion that pericytes do not contribute to fibrotic scar tissue after brain injury^30^. While our cortical stab wound results are in agreement with their findings, the contribution of Tbx18-expressing perivascular cells to CNS fibrosis should be reassessed in a lesion model that generates substantial fibrotic scar tissue.

MCAO-induced hypoxic lesions restricted to the striatum showed an increase of type A pericyte-derived PDGFRβ^+^ stromal cells in association with the vascular wall, but little to no stromal cells outside the vessel wall (Fig. 4 and Fig. 7). In sharp contrast, a large number of type A pericyte-derived PDGFRβ^+^ stromal cells could be found outside the vessel wall after cortico-striatal ischemic lesions (Fig. 7 and Supplementary Fig. 5). These observations may explain previous results regarding pericyte detachment from the vascular wall^31,32,33^ and suggest that the location and magnitude of the ischemic insult dictate the distribution of type A pericyte progeny and generation of stromal fibroblasts. In the lesion core of subacute/chronic ischemic stroke lesions in humans, we found nearly all PDGFRβ^+^ stromal cells associated with the vascular wall (Fig. 6g, h).

In the GL261 murine glioma model, the vasculature is known to present low coverage by pericytes^22^, including type A pericytes. However, despite their reduced number, we observed PDGFRβ^+^ stromal cells outside tumor vessels interspaced with glioma cells, partially derived from type A pericytes (Fig. 5 and Fig. 7). In human GBM tumors, most PDGFRβ^+^ cells remained in close association with the stroma vasculature (Fig. 6i,j).

Pericytes have also been shown to be the main contributors to ECM-producing (myo)fibroblasts observed in dermal, renal, hepatic and pulmonary fibrotic tissue^34–37^, suggesting functional similarities among pericytes in the CNS and peripheral organs. Nonetheless, while pericyte heterogeneity has not been addressed in the context of peripheral organ fibrosis, we define the origin of PDGFRβ-expressing stromal cells to a specific perivascular subset, only accounting for 10% of all CNS pericytes. This distinction is of special importance for the design of potential therapeutic strategies to improve axonal regeneration and functional recovery following CNS injury^6^, as it may allow specific targeting of fibrotic pericytes without compromising the vast majority of pericytes and their functions (*i.e*. maintenance of the blood-brain/spinal cord barrier).

## Methods

### Transgenic Mice

GLAST-CreER^T2^ transgenic mice^12^ were crossed to the Rosa26-enhanced yellow fluorescent protein (EYFP) Cre-reporter line^38^ (obtained from the Jackson Laboratory, B6.129X1-Gt(Rosa)26Sor^tm1(EYFP)Cos^/J, JAX stock: 006148) to generate GLAST-CreER^T2^;R26R-EYFP, in which CreER^T2^ is hemizygous and Rosa26-EYFP is either heterozygous or homozygous. All animals were in a C57BL/6J genetic background and ≥ 8 weeks old at the onset of experiments.

Animals were housed in group in a pathogen-free facility with controlled humidity and temperature, with 12:12-hour light:dark cycles and free access to food and water. All experimental procedures were performed in accordance to the Swedish and European Union guidelines and approved by the institutional ethical committees (*Stockholms Norra Djurförsöksetiska Nämnd* or *Malmö/Lunds Djurförsöksetiska Nämnd*).

### Genetic labeling of transgenic mice

Genetic recombination in GLAST-CreER^T2^;R26R-EYFP animals was induced by a daily intraperitoneal injection of 2 mg of tamoxifen (Sigma, 20 mg/ml in 1:9 ethanol:corn oil) for five consecutive days, as previously^3,6^.

### Clearing period following tamoxifen-induced genetic recombination

A potential confound with tamoxifen-inducible mouse studies is the effect of residual tamoxifen in the CNS at the time of injury. Tamoxifen and its active metabolite 4-hydroxytamoxifen have a half-life of 6–12 h in the mouse^39^. Analysis of CreER^T2^ distribution in the adult mouse spinal cord 6 days after the last tamoxifen administration has demonstrated that there is no CreER^T2^ in the nucleus of cells at this time, directly demonstrating that tamoxifen has been cleared at this time point^15^. Moreover, recent studies in the mouse spinal cord and brain have reinforced that 7 days is enough time for tamoxifen to be fully metabolized to ineffective concentrations that are insufficient to result in further recombination^40,41^.

Therefore, in this study, injuries (spinal cord injuries, stab wound and ischemic stroke lesions) and tumor induction (GL261 murine glioma model) were performed after a 7-days clearing period following the last tamoxifen injection. Experimental autoimmune encephalomyelitis (EAE) was induced following a 50-day clearing period after the last tamoxifen injection, because the clinical severity of the disease was diminished if MOG-peptide immunization took place within 10 days after tamoxifen treatment^18^.

### Surgical procedures

For all surgical procedures, animals were anesthetized with 4% isoflurane until unconscious followed by 2% isoflurane during surgery. All animals received analgesia (Temgesic/Buprenorphine, Schering-Plough, 0.1 mg/kg body weight and Rimadyl/Carprofen, Pfizer, 5 mg/kg body weight; subcutaneous injection) for postoperative pain relief and a uniform layer of eye gel (Viscotears/Carbomer, 2 mg/g, Novartis) was applied onto the eye ball to prevent drying. All animals received local anesthesia (Xylocaine/Lidocaine, AstraZeneca, 10 mg/ml and Marcain/Bupivacaine, AstraZeneca, 2mg/kg body weight; 2 drops on the spinal cord surface) 2-3 min prior to craniotomies and spinal or brain lesions. For recovery from surgery, animals were placed on a heating pad and only returned to their home cage once they were fully awake.

#### - Penetrating spinal cord injury: Dorsal *funiculus* incision model

A laminectomy was performed at the mid-thoracic level to expose the dorsal portion of the spinal cord and the dorsal *funiculus* and adjacent grey matter were cut transversely to a depth of 0.6 mm with microsurgical scissors. This incision was extended rostrally with microsurgical scissors to span one spinal segment^6,15^. Both male and female mice were used.

Animals were sacrificed at 5 days (n=3), 14 days (n=6) and 6 weeks (n=4) after SCI. A group of recombined GLAST-CreER^T2^;R26R-EYFP (n=3) mice received no injury and served as uninjured control.

#### - Non-penetrating spinal cord injury: Complete spinal cord crush model

A laminectomy was performed to expose the dorsal portion of the spinal cord at the thoracic level 10 (T10). The spinal cord was then fully crushed for 2 seconds with Dumont No.5 forceps (11295-00, Fine Science Tools) without spacers and that had been filed to a width of 0.1 mm for the last 5 mm of the tips^16,42^. Since this injury paradigm minimizes mechanical disruption of the *dura mater*, it limits confounding factors such as invasion of *dura*-derived meningeal fibroblasts into the lesion.

To replace blood loss and prevent/treat bladder infections and postoperative pain, antibiotics (Trimetoprimsulfametoxazol/Tribrissen vet, 400 mg/ml Sulfadizin, 80 mg/ml Trimetoprim, MSD Animal Health, 100 mg/kg body weight/24hr; subcutaneous injection), sterile Ringer’s solution (500 ul; subcutaneous injection) and analgesics (Rimadyl/Carprofen, Pfizer, 5 mg/kg body weight, once a day and Temgesic/Buprenorphine, Schering-Plough, 0.1 mg/kg body weight, twice per day; subcutaneous injection), were administered in the first 3 days following surgery. Bladders were manually expressed 2-3 times per day until the end of the experiment. Only female mice were used for this severe SCI model.

Animals were sacrificed at 5 days (n=4), 14 days (n=5) and 7 weeks (n=3) after SCI. A group of recombined GLAST-CreER^T2^;R26R-EYFP (n=3) mice received no injury and served as uninjured control.

#### - Stab lesions

Animals were placed in a stereotaxic frame and a unilateral craniotomy (diameter of ∼3 mm) was performed between cranial sutures bregma and lambda above the right cerebral cortex (adapted from^17^). An incision wound was introduced into the right cortical parenchyma by inserting a sterile surgical scalpel blade (Kiato, #11) at 2.0 mm lateral and 1 mm anterior to bregma. To produce stab wound lesions restricted to the cerebral cortex, the scalpel blade was lowered −0.5 mm from the brain surface in the dorsoventral axis to avoid direct damage to the deepest cortical layers and underneath white matter, minimizing neuroblast migration from the subventricular zone of the lateral ventricles and subgranular zone in the dentate gyrus of the hippocampus into the injured cortex. For cortico-striatal stab wound lesions, the blade was lowered −3 mm from the brain surface in the dorsoventral axis.

The lesions were created by moving the blade over 1.3 mm back and forth along the anterior-posterior axis from 1 mm anterior to 0.3 mm posterior to bregma. The craniotomy was then covered with bone wax (Ethicon) in order to preserve the meninges and the brain surface. Both male and female mice were used. Animals were sacrificed at 5 days (n=3), 14 days (n=4) and 7 weeks (n=3) after cortico-striatal stab lesions. The contralateral side to the lesion served as control (n=10). A separate cohort of animals was sacrificed at 14 days after stab lesions restricted to the cerebral cortex (n=3).

#### - Middle cerebral artery occlusion (MCAO)

Intraluminal filament model of middle cerebral artery occlusion was used to induce stroke as described^13,43^. Briefly, following a sagittal midline incision on the ventral surface of the neck, the common carotid artery (CCA) and its proximal branches were isolated. After the CCA and the external carotid artery (ECA) were ligated using a 4-0 silk suture (Ethicon), the internal carotid artery (ICA) was temporarily clipped with a metal microvessel clip. A small incision was made on the anterior surface of the ECA to insert a silicon-coated nylon microfilament (size 7.0) The microfilament was advanced through the ICA until the branch point of the MCA and secured using the suture around the ECA. The filament was carefully removed 35 min after the occlusion to restore the blood flow in the MCA and the ECA was ligated permanently. Only male mice were used.

Animals were sacrificed at 5 days (n=3), 14 days (n=3) and 7 weeks (n=3) after stroke induction. The contralateral side to the lesion served as control (n=9). A separate cohort of animals presented ischemic lesions that, in addition to the striatum, extended into the cerebral cortex (cortico-striatal stroke), and were analysed separately (n=2; 5 dpi).

#### - Experimental Autoimmune Encephalomyelitis (EAE)

To induce EAE, animals were anesthetized with isoflurane and immunized subcutaneously at the dorsal tail base with 300 µg of rodent myelin oligodendrocyte glycoprotein (MOG) peptide (aminoacids 35–55) emulsified in Complete Freund’s Adjuvant (CFA) containing 5 mg/ml inactivated Mycobacterium tuberculosis (Difco). *Pertussis* toxin (75 ng/mouse in PBS) was injected intravenously on the day of immunization and 48 hours later.

Animals allocated to control groups received *Pertussis* toxin only or Complete Freund’s Adjuvant (CFA) containing 5 mg/ml inactivated *Mycobacterium tuberculosis* plus *Pertussis* toxin, but no MOG peptide.

The animals were weighted and neurological deficits assessed daily until the end of the experiment according to previous scoring methods^44,45^: 0 – no apparent symptoms, 1-flaccid tail or wobbling gait, 2 – flaccid tail and wobbling gait, 2.5 – single limb paresis and ataxia, 3 – double limb paresis, 3.5 – single limb paralysis and paresis of second limb, 4 – complete paralysis of hind- and fore-limbs, 4.5-moribund, 5-dead. Both male and female mice were used. Animals were sacrificed 30 days post immunization with MOG peptide in CFA (EAE group, n=4), or *Pertussis* toxin alone and/or CFA (control group, n=5). Only animals that had reached a clinical score of at least 3 by the time of sacrifice and presented chronic neurological deficits and not relapse-remitting symptoms were analyzed in the EAE group.

#### - Syngenic mouse model of glioblastoma

GL261 mouse glioma cells (DSMZ, ACC 802) were maintained in DMEM 1X, high glucose, GlutaMAX (+pyruvate) medium (Life Technologies, 31966-021) supplemented with 10% heat inactivated Fetal Bovine Serum and 1x Pen-Strep at 37°C in a humidified atmosphere containing 5% CO_2_. Cells were cultured in the absence of antibiotics before transplantation.

Animals were head-fixed in a stereotaxic frame and a burr hole was drilled over the sensorimotor cortex to expose the brain. Fifty thousand GL261 cells were resuspended in 1 μl of sterile phosphate buffered saline (PBS) and injected in the striatum, at 2.3 mm lateral to the midline, 0.1 mm posterior of the bregma and 2.3 mm ventral from dura, at a rate of 0.1 μl/min using a 10 μl syringe with a 26-gauge beveled needle tip (Hamilton, 901RN) and a microinjector (UltraMicroPump III and Micro4 microsyringe pump controller, World Precision Instruments). After injection, the needle was kept in place for an additional 5 min to allow the cells to diffuse and prevent backflow of the cells to the surface, and then slowly withdrawn. Both male and female mice were used. Animals (n=4) were sacrificed 3 weeks after inoculation of the tumor cells. The contralateral side to the lesion served as control.

### Human tissue collection and ethical compliance

This study complied with all relevant ethical regulations regarding experiments involving human tissue samples. Ethical permission for this study was granted by the Regional Ethics Committee of Sweden (2010/313-31/3).

Supplementary tables 1-4 show the clinical and neuropathological data of all subjects included in the study.

#### - Stroke and glioblastoma samples

The institutional review boards and the local ethics committee of the Medical Faculty of the University of Erlangen-Nuremberg, Erlangen, Germany, approved the study (issued ethical votes No. 4821, 104_13B and 331_14B) and informed consent was obtained from the relatives of all analyzed patients. For stroke samples, tissue collection was performed in such cases that had a history of a supratentorial territorial or lacunar stroke involving cerebral cortical and subcortical areas and died from a non-neurological cause. The lesion was cut out according to the macroscopically visible infarct borders including a 0.5-cm rim of surrounding healthy tissue. Healthy occipital cortex and healthy tissue from the contralateral hemisphere in corresponding topography served as control tissue. Brain tissue was frozen and stored at −80 °C until further analysis. For glioblastoma samples, initial histological analysis for verification of tumor type and WHO classification^46^ was performed by local experienced neuropathologists on formalin embedded tissue pieces (frozen tumor tissue samples, 20 µm microtome sectioning and fixed in 4% formaldehyde buffered in PBS for 30 min). Standard hematoxylin and eosin staining, and immunohistochemical analyses for GFAP, MAP2 and pan cytokeratin-1 (KL-1) confirmed tumor types (glioblastoma, WHO grade IV). Brain tissue was frozen and stored at −80 °C until further analysis.

#### - Multiple sclerosis samples

Multiple sclerosis tissue samples and associated clinical and neuropathological data were supplied by the UK Multiple Sclerosis Tissue Bank, supported by the Multiple Sclerosis Society of Great Britain and Northern Ireland, in partnership with Imperial College London. The ethical permission was granted by the Regional Ethics Committee for UK (Research Ethics Committee for Wales, 08/MRE09/31+5). All samples have been donated with informed consent for use in future research. Spinal cord tissue was frozen and stored at −80°C until analysis.

#### - Spinal cord injury samples

Human spinal cord injury samples and related clinical and neuropathologial information were obtained from the International Spinal Cord Injury Biobank (ISCIB), which is housed in Vancouver, BC, Canada. Permission for *post-mortem* spinal cord acquisition and for sharing of biospecimens was granted by the Clinical Research Ethics Board (CREB) of the University of British Columbia, Vancouver, Canada (Ethics certificate of full board approval H19-00690). All biospecimens were collected from consented participants or their next-of-kin. Provided formalin-fixed and paraffin-embedded tissue sections were stored at 4°C until analysis.

### Histology and Immunohistochemistry

Animals were euthanized by intraperitoneal injection of an overdose of sodium pentobarbital and transcardially perfused with cold PBS followed by 4% formaldehyde in PBS. Brains and spinal cords were dissected out and post-fixed in 4% formaldehyde in PBS overnight at 4^°^C. Spinal cords were then cryoprotected in 30% sucrose and coronal or sagittal cryosections were collected on alternating slides. Coronal and sagittal brain sections were obtained using a vibratome.

For human glioblastoma, stroke and multiple sclerosis tissue samples, fresh frozen sections were prepared with a cryostat (14-20 μm) and post-fixed in PBS-buffered 2% formaldehyde (wt/vol) for 10 min according to standard procedures.

Formalin-fixed and paraffin-embedded coronal spinal cord tissue sections (5 μm thick) were obtained from spinal cord injury patients. For immunohistochemistry, spinal cord sections were deparaffinized in xylene and rehydrated in a descending ethanol series. Antigen retrieval was performed in citraconic acid solution (pH = 7.4; 0.05% citraconic acid) for 20 min in a domestic steamer^47^. The sections were allowed to cool down for 20 min before immunostaining was started.

Sections were incubated with blocking solution (10% normal donkey serum in PBS, with 0.3% Triton X-100) for 1 hour at room temperature, and then incubated at room temperature overnight in a humidified chamber with primary antibodies diluted in 10% normal donkey serum. The following primary antibodies were used to immunostain mouse tissue: CD31 (1:100, rat, BD Biosciences), GFAP (1:200, guinea pig, Synaptic Systems; 1:1000, mouse directly conjugated to Cy3, Sigma-Aldrich; 1:1000, chicken, Millipore), GFP (1:2000, goat directly conjugated to FITC, Abcam; 1:10000, chicken, Aves Labs), PDGFRβ (1:200, rabbit, Abcam) and Podocalyxin (1:200, goat, R&D Systems). An antibody against NG2 (1:200, rabbit, Millipore) was used to detect GL261 tumor cells^48^. The following primary antibodies were used to immunostain human tissue: PDGFRβ (1:100, goat, R&D Systems), GFAP (1:200, rabbit, Dako; 1:1000, chicken, Millipore), MBP (1:500, rat, AbD Serotec; 1:1000, mouse, Covance), CD31 (1:100, mouse, Dako), Podocalyxin (1:200, goat, R&D Systems), von Willebrand factor (1:200, rabbit, Dako) and UEA-1 (1:200, directly conjugated to FITC or Rhodamine, Vector Labs). Following primary antibody incubation and washing, antibody staining was revealed using species-specific fluorophore-conjugated (Alexa Fluor 488, Cy3, Alexa Fluor 647 from Jackson Immunoresearch) or biotin-conjugated secondary antibodies (1:500, Jackson Immunoresearch). Biotinylated secondary antibodies were revealed with fluorophore-conjugated streptavidin (1:500, Cy3 from Jackson Immunoresearch). Control sections were stained with secondary antibody alone. Cell nuclei were stained with 4’,6’-diamidino-2-phenylindole dihydrochloride (DAPI, 1 µg/ml, Sigma-Aldrich). Sections were coverslipped using Vectashield Antifade Mounting Medium (Vector Labs, H-1000).

For hematoxylin & eosin (H&E) staining, tissue sections were washed in ddH2O for 5 min, followed by 1 min immersion in hematoxylin (Vector Labs, H-3401) and washed again under tap water until clear. Following the immersion in ddH2O, sections were stained with eosin (Vector Labs, H-3502) for 3 min. Sections were finally washed in ddH2O for 1 minute, dehydrated and coverslipped with VectaMount permanent mounting media (Vector Labs, H-5000).

### Imaging and Quantitative Analysis

Images were acquired with a Leica CTR6000 bright-field microscope, a Zeiss Axioplan 2 upright epifluorescent microscope or a Leica TCS SP8X confocal microscope. Image processing and assembly were performed with ImageJ/Fiji (version 2.0.0-rc-43/1.51j for Mac), Adobe Photoshop CC 2018 19.1.1 release and Illustrator CC 2018 22.0.1 release for Mac.

Coronal (dorsal *funiculus* incision SCI model) or sagittal (complete crush SCI model) spinal cord sections, spanning the injury site and 0.4 mm rostral and caudal to the injury site, were collected on 20 alternating slides at 20 μm thickness and used for quantifications. Matched segments of uninjured spinal cords were sectioned and collected in a similar fashion. For EAE spinal cord samples, 20 μm thick coronal sections covering cervical, thoracic and lumbar spinal segments were collected on 50 alternating slides and used for quantifications. Control spinal cord samples (*Pertussis* toxin alone and/or CFA) were sectioned and collected in a similar fashion. Brains with stab wound injury were sectioned at 30 μm thickness and collected in 20 alternating series. Coronal sections (between rostrocaudal levels 1.0 mm anterior to 0.7 mm posterior of bregma) were used for quantification and sagittal sections used for the images presented in Figure 2. For MCAO and mouse glioma samples, coronal brain sections covering the ischemic stroke core and tumor core (between rostrocaudal levels 1.0 mm anterior to 0.7 mm posterior of bregma), respectively, were collected in 20 alternating series at 30 μm thickness and used for analysis.

All quantifications were done in 3 alternate sections per animal covering the lesion epicenter and spaced 400 μm apart. The contralateral side to the lesion served as control and was used for mouse glioma, stroke and stab wound control analyses. Uninjured spinal cord samples were used as controls for spinal cord injury quantifications. Three alternate sections per animal, spaced 400 μm apart were sampled for control analyses. For EAE and respective control (pertussis toxin alone and/or CFA) samples, 3 cervical, 3 thoracic and 3 lumbar sections per animal, spaced 1000 μm apart were analysed.

To determine the lesion core area, sections were immunostained for PDGFRβ, GFAP and DAPI. The lesions were imaged using a Zeiss Axioplan 2 epifluorescent microscope and the lesion core area, defined as the GFAP negative area within the GFAP-defined borders filled with PDGFRβ-expressing stromal cells, was manually outlined. For EAE samples, the area covered by the multiple scar-like clusters of PDGFRβ-expressing stromal cells and reactive astrocytes was manually delineated. Area measurements were carried out using the ImageJ/Fiji software (version 2.0.0-rc-43/1.51j for Mac).

The total number of type A-pericyte derived stromal cells, defined as EYFP-positive cells co-expressing PDGFRβ within the lesion core, was assessed in spinal cord or brain sections immunolabeled for GFP (that cross-reacts and recognizes EYFP), PDGFRβ, GFAP and DAPI. Similarly, matched areas on the striata contralateral to the lesion side or uninjured/control spinal cord sections were used to quantify the total number of type A pericytes (EYFP-positive cells co-expressing PDGFRβ enwrapping the blood vessel wall) under control conditions. The data were averaged and presented as total number of recombined (EYFP^+^) PDGFRβ-expressing cells per section.

The density of type A pericyte-derived stromal cells at the lesion epicenter, a measure that reflects cell clustering, was calculated by dividing the total number of recombined PDGFRβ-expressing cells per section by the corresponding lesion core area. For uninjured/control spinal cord quantifications, the total number of type A pericytes per spinal section was divided by the area of the corresponding spinal cord section. For control brain quantifications, the total number of type A pericytes present in the striatum contralateral to the lesion side was divided by the corresponding striatal area. The data were pooled and presented as the mean number of recombined PDGFRβ-expressing cells per area.

To determine the proportion of recombined cells either associated with or dissociated from the blood vessel wall within the lesion core, tissue sections were immunostained for the endothelial markers CD31 or podocalyxin, GFP, PDGFRβ and DAPI. Four randomly selected fields (0.83mm × 0.66 mm) per section were photographed and used for analysis. The number of recombined PDGFRβ-positive cells in contact with the endothelium (ON vessel) or away from the blood vessel wall (OFF vessel) was manually counted and the data expressed as a percentage of total recombined PDGFRβ-expressing cells. The percentage of type A pericytes ON or OFF vessels under uninjured/control conditions was assessed in a similar fashion.

To assess the proportion of PDGFRβ-expressing stromal cells that originates from type A pericytes, the total number of recombined (EYFP^+^) and non-recombined (EYFP^−^) PDGFRβ-expressing stromal cells within the lesion core was manually counted and used to calculate the percentage of recombined PDGFRβ-expressing cells out of total PDGFRβ-expressing cells. Likewise, uninjured/control sections were used to calculate the percentage of type A pericytes out of all PDGFRβ-expressing pericytes.

For EAE samples, the number of lesions per section was quantified in 3 cervical, 3 thoracic and 3 lumbar sections per animal spaced 1000 um apart. A lesion was identified as a large aggregate of PDGFRβ-expressing stromal cells and reactive astrocytes. The anatomical location of the lesion (dorsal, intermediate, ventrolateral or ventral white matter) was noted and the results were presented as number of lesions per section at the cervical, thoracic or lumbar spinal levels.

### Statistical Analysis

No statistical methods have been used to predetermine sample size (sample sizes were determined based on previous experience). Data are presented as mean ± standard error of the mean (s.e.m.). Individual data points are plotted for some graphs. Sample sizes (*n*) of animals, number of biological repeats of experiments and statistical methods used are indicated in the corresponding figure legends. Sample groups were too small to test for Gaussian distribution (<8 samples) and parametric tests were preferred because nonparametric tests lack suitable power to detect differences in small sample groups. *P* values were calculated using two-tailed unpaired Student’s *t*-test, one way- or two-way ANOVA. Repeated measures were applied when appropriate. *Post hoc* correction tests were employed following significance with an ANOVA and described in the respective figure legend.

Differences were considered statistically significant at P values below 0.05. All data were analyzed using GraphPad Prism version 6.0g software for Mac.

## Data Availability

The authors declare that all data supporting the findings of this study are included in this published article (and its supplementary information files). Source Data for Figs. 1–5 and Supplementary Fig. 3 are provided with the paper.

## Acknowledgments

We thank Göritz lab members for valuable comments on the manuscript. D.O.D. was supported by the Foundation for Science and Technology from the Portuguese government (SFRH/BD/63164/2009). H.B.H. was supported by a grant from the German Research Foundation (Hu1961/2-1). C.G. is a Hållsten Academy and a Knut and Alice Wallenberg Academy Fellow. Research in the C.G. lab was supported by the *European Union’s Seventh Framework Programme (FP7)/*ERC-2012-StG 310938 PERICYTESCAR, Swedish Research Council, Swedish Brain Foundation, Ming Wai Lau Centre, JPND DACAPO-AD and Wings for Life Foundation. Multiple sclerosis tissue samples and associated patient clinical and neuropathological data were supplied by the UK Multiple Sclerosis Tissue Bank, supported by the Multiple Sclerosis Society of Great Britain and Northern Ireland, registered charity 207495. We also thank Dr. Brian Kwon and the International Spinal Cord Injury Biobank (ISCIB) located at the University of British Columbia in Vancouver, BC, Canada for providing tissue sections from spinal cord injured patients for histologic analysis.

## Author Contributions

D.O.D., J.K., Y.K., C.P.E., J.T. and A.E. performed experiments and/or analyses. H.B.H. and L.B. supplied human tissue. D.O.D. and C.G. designed experiments. L.B., Z.K., O.L., H.B.H., J.F. and C.G. supervised experiments. C.G. coordinated the study. D.O.D. and C.G. wrote the manuscript.

## Declaration of Interests

The authors declare no competing financial interests.

**Correspondence and requests for materials** should be addressed to C.G.

