## Supplementary material for "Pericyte-derived fibrotic scarring is conserved across diverse central nervous system lesions"

Dias, D.O. *et al.*

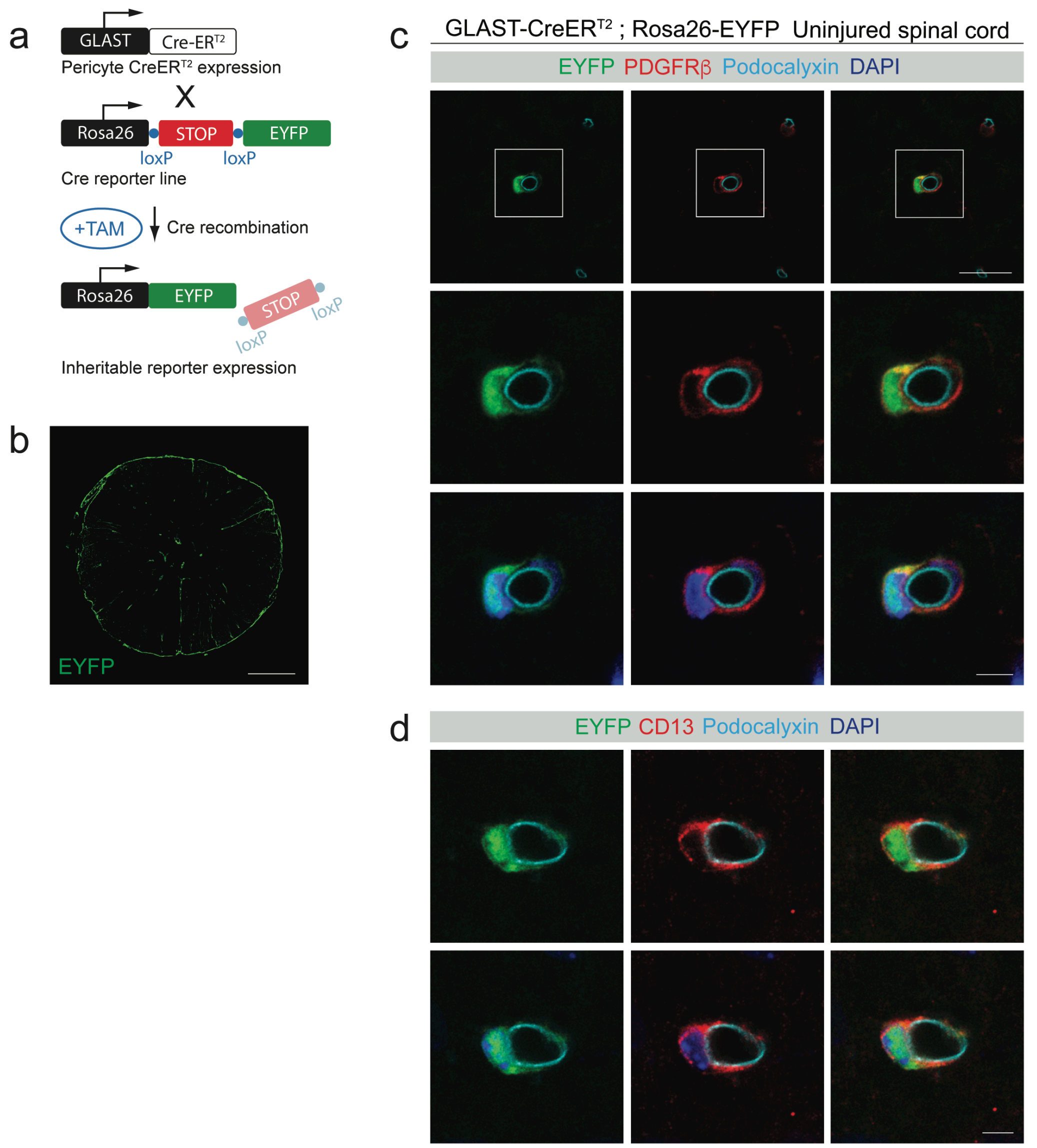


Supplementary Figure 1

Supplementary Figure 1 | **Genetic labeling of type A pericytes in the adult spinal cord**

(**a**) Schematic depiction of the strategy to induce genetic recombination and labeling of a subset of perivascular cells, named type A pericytes. Type A pericytes (expressing the GLAST-CreER$^{T2}$ transgene) undergo tamoxifen-mediated genetic recombination and turn on EYFP expression. Type A pericytes and progeny can be traced by stable and inheritable labeling with EYFP.

(**b**) Distribution of recombined cells (EYFP$^{+}$) in the adult uninjured spinal cord of GLAST-CreER$^{T2}$; R26R-EYFP mice.

(**c,d**) Type A pericytes (EYFP$^{+}$) encapsulate the endothelial tube (podocalyxin) and express the pan pericyte markers PDGFRβ (c) and CD13 (d) in the adult uninjured mouse spinal cord.

Scale bars show 400 μm (b), 20 μm (c,d) and 5 μm (close ups in c,d). Nuclei are labeled with 4′,6-diamidino-2-phenylindole (DAPI). All images show coronal sections.

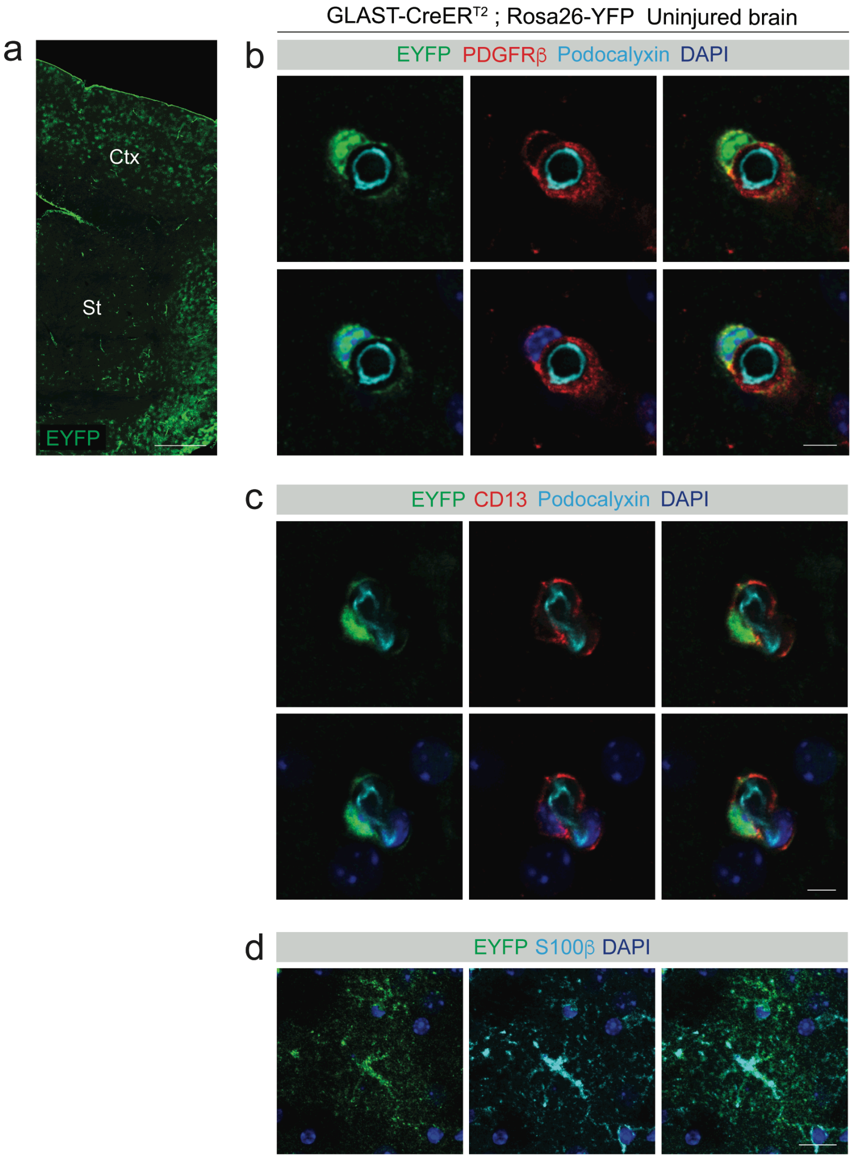


Supplementary Figure 2

Supplementary Figure 2 | **Genetic labeling of type A pericytes in the adult brain**

(**a**) Distribution of recombined cells ($EYFP^+$) in the adult uninjured forebrain of $GLAST\text{-}CreER^{T2}$; R26R-EYFP mice.

(**b**,**c**) Type A pericytes ($EYFP^+$) encapsulate the endothelial tube marked with an antibody against podocalyxin and express PDGFRβ (b) and CD13 (c).

(**d**) In addition to type A pericytes (b, c), recombination ($EYFP^+$) occurs in parenchymal astrocytes expressing S100β.

Scale bars show 600 μm (a), 20 μm (d) and 5 μm (b,c). Nuclei are labeled with 4′,6-diamidino-2-phenylindole (DAPI). a-d show coronal sections.

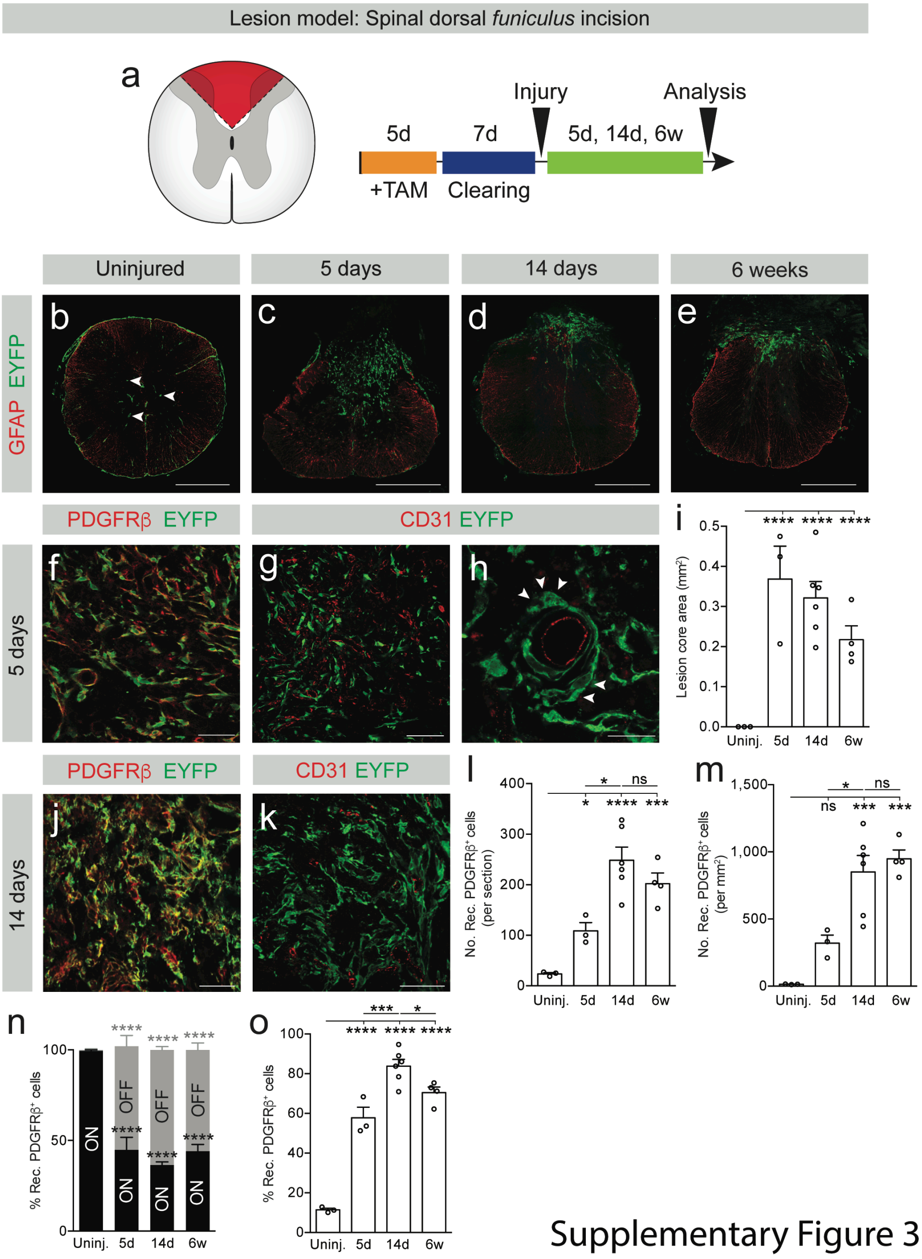


Supplementary Figure 3

Supplementary Figure 3 | **Type A pericyte progeny is the main source of stromal fibroblasts that form fibrotic scar tissue after penetrating spinal crush injury**

(**a**) Dorsal *funiculus* incision spinal cord injury model and experimental timeline.

(**b**-**e**) Distribution of recombined cells ($EYFP^+$) in the uninjured spinal cord (b) and at 5 days (c), 14 days (d) and 6 weeks (e) following spinal cord injury. Arrows in (b) point at examples of $EYFP^+$ type A pericytes.

(**f**) Recombined cells co-express the stromal marker PDGFRβ, 5 days after injury.

(**g**,**h**) A fraction of recombined cells is located outside the vascular wall ($CD31^{+}$) at 5 days post-injury. Arrowheads in (h) point at examples of recombined cells that are found in distance to the vascular wall.

(**i**) The lesion core decreases in size, as the scar condenses.

(**j**,**k**) A fraction of recombined $PDGFR\beta^{+}$ cells (j) remains outside the blood vessel wall (k) at 14 days after injury.

(**l**,**m**) The number (l) and density (m) of recombined $PDGFR\beta^{+}$ cells peak at 14 days post-injury and remain comparable thereafter.

(**n**) Afetr injury a fraction of recombined $PDGFR\beta^{+}$ cells are located outside of the vessel wall (OFF vessel).

(**o**) Following injury, type A pericyte progeny are the main contributors to PDGFRβ-expressing stromal cells populating the lesion core.

Scale bars show 500 μm (b-e), 100 μm (G), 50 μm (f, j, k) and 20 μm (h). Data shown as mean ± s.e.m. n=3 (Uninjured), n=3 (5 dpi), n=6 (14 dpi), n=4 (6 wpi). ns, non-significant; *$P<0.05$, ***$P<0.001$, ****$P<0.0001$ by One-Way ANOVA followed by Holm-Sidak *post hoc* test. All images show coronal sections.

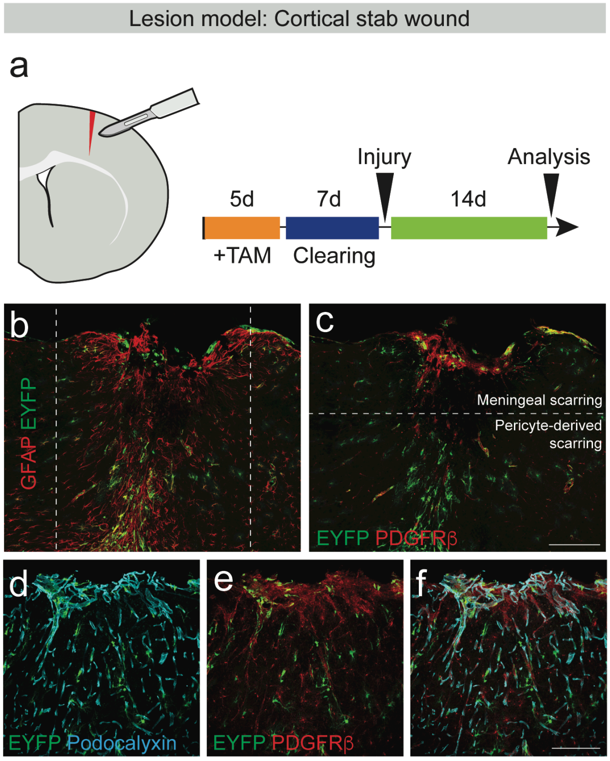


Supplementary Figure 4

Supplementary Figure 4 | **Stab lesions restricted to the cerebral cortex do not generate extensive fibrotic scarring**

(**a**) Cortical stab lesion model and experimental timeline.

(**b**,**c**) Stab lesions restricted to the cerebral cortex induce widespread gliosis ($GFAP^+$; b) and generate meningeal-derived fibrotic scarring close to the brain surface ($PDGFR\beta^+$), but do not trigger extensive pericyte-derived fibrotic scarring, as observed by limited $EYFP^+$/$PDGFR\beta^+$ cells at 14 dpi (c).

(**d-f**) Recombined cells ($EYFP^+$) do not leave the blood vessel wall (podocalyxin) following cortical stab lesions.

Scale bars represent 200 μm (d-f) and 250 μm (b, c). (b,c) and (d-f) show paired images. All images show sagittal sections.

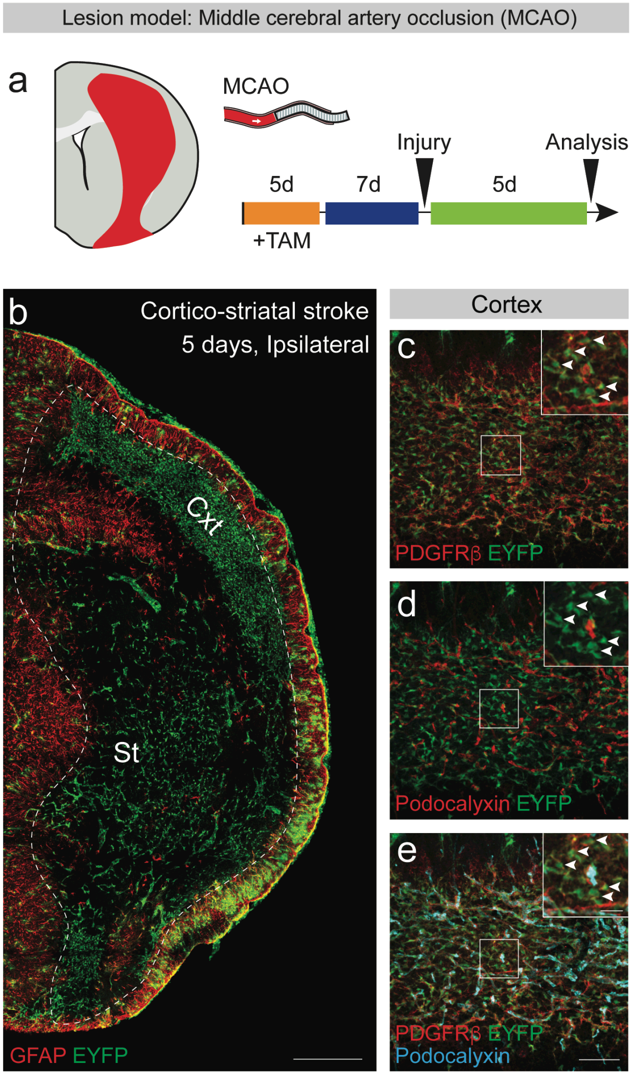


Supplementary Figure 5

Supplementary Figure 5 | **Type A pericyte-derived cells contribute to stromal fibroblasts after cortico-striatal ischemic stroke**

(**a**) Middle cerebral artery occlusion (MCAO) stroke model inducing cortico-striatal ischemic lesions and experimental timeline.

(**b**) Distribution of recombined (EYFP$^+$) cells at 5 days following injury. Partially recombined reactive glial cells (GFAP$^+$) surround the ischemic lesion core (dashed line), filled with EYFP$^+$ stromal cells.

(**c-e**) Recombined cells in the ischemic stroke core at 5 days post-injury co-express the stromal marker PDGFRβ (c) and are found at a distance to the vascular wall (d) in the cortex (arrowheads). (e) shows merged images of (c) and (d). Insets show a higher magnification of boxed areas.

Scale bars represent 500 μm (b), 100 μm (c-e) and 25 μm (Insets c-e). (c, d and e) represent paired images. All images show coronal sections.

Supplementary table 1 | **Clinical and neuropathological data of Spinal Cord Injury patients**

| Case ID | Age | Sex | Injury level* | AIS grade | Injury-death interval (days) | Tissue harvest *post mortem* (hrs) | Type of Injury# | Diagnosis |
|---|---|---|---|---|---|---|---|---|
| BB1 | 55 | M | C6 | A | 10 | 26 | MC | Three column fracture dislocation |
| BB3 | 80 | F | C6 | B | 17 | 49 | MC | Bilateral facet dislocation |
| BB4 | 76 | M | C7 | A | 34 | 14 | MC | Three column fracture dislocation |
| BB5 | 82 | M | C6 | A | 9 | 13 | MC | Bilateral facet dislocation |
| BB9 | 95 | M | C4 | C | 16 | 26 | C/C | Hyperextension, avulsion flakes, traumatic disc |
| BB6 | 83 | M | C4 | D | 61 | 15 | C/C | Hyperextension, avulsion flakes, traumatic disc |

* Cervical ; Number indicates vertebrae
### MC – Massive compression type injury ; C/C – Contusion/Cyst type injury

Supplementary table 2 | **Clinical and neuropathological data of Multiple Sclerosis patients**

| Case ID | Age | Sex | Disease type* | Disease duration (years) | Lesion stage‡ | Tissue harvest *post mortem* (hrs) | Diagnosis |
|---|---|---|---|---|---|---|---|
| MS 058 | 51 | F | SPMS | 21 | AL / CAL | 15 | Chronic MS |
| MS 062 | 49 | F | SPMS | 19 | AL / CAL | 10 | Chronic MS |
| MS 066 | 86 | F | ND | 56 | CAL | 21 | Chronic MS |
| MS 074 | 64 | F | SPMS | 36 | CAL | 7 | Chronic MS ; Significant demyelination |
| MS 092 | 37 | F | SPMS | 17 | CAL | 26 | Chronic MS ; Epileptic seizures |
| MS 097 | 55 | M | SPMS | 22 | CAL | 31 | Chronic MS |
| C14 | 64 | M | NA | NA | NA | 18 | Control tissue (spinal cord looks normal) |
| C37 | 84 | M | NA | NA | NA | 5 | Control tissue (spinal cord looks normal) |
| C39 | 82 | M | NA | NA | NA | 21 | Control tissue (spinal cord looks normal) |
| C43 | 87 | F | NA | NA | NA | 12 | Control tissue (spinal cord looks normal) |

* SPMS – Secondary Progressive Multiple Sclerosis ; ND – Not documented (little medical history available)
‡ AL – Active Lesion ; CAL – Chronic Active Lesion
NA – Non applicable

Supplementary table 3 | **Clinical and neuropathological data of Stroke patients**

| Case ID | Age | Sex | Type of stroke | Location of stroke | Size of infarction ($cm^3$) | Stroke-death interval | Tissue harvest *post mortem* (hrs) | Cause of infarction |
|---|---|---|---|---|---|---|---|---|
| 16-573 | 65 | M | Territorial (Arteria cerebri media) | Right frontal cortex | ≈ 36 | 3 months | 36 | Atrial fibrillation (cardiac source) |
| 14-24 | 83 | M | Territorial (Arteria cerebri media) | Right Basal ganglia (including thalamus) | ≈ 26 | 7 weeks | 24 | Stenosis of Internal carotid artery (NASCET 80%*) |
| 14-16 | 86 | F | Lacunar | Left striatum | ≈ 2 | 4 weeks | 36 | Artherothrombotic |
| 13-69 | 81 | F | (i) Lacunar ; (ii) Territorial (Arteria cerebri media) | (i) Left striatum (ii) Left parietal cortex | (i) ≈ 2 (ii) ≈ 48 | (i) 31 days (ii) 26 months | 36 | Atrial fibrillation (cardiac source) |

*NASCET – North American Symptomatic Carotid Endarterectomy Trial

Supplementary table 4 | **Clinical and neuropathological data of Glioblastoma patients**

| Case ID | Age at operation | Sex | Type of tumor | Location of tumor | Tumor size (cc) |
|---|---|---|---|---|---|
| FP190138 | 76 | F | Glioblastoma (WHO Grade IV; Astrocytoma) | Left temporal lobe | ≈ 54 |
| FP220453 | 64 | F | Glioblastoma (WHO Grade IV; Astrocytoma) | Left fronto-temporal lobe | ≈ 60 |
| FP070741 | 75 | F | Glioblastoma (WHO Grade IV; Astrocytoma) | Corpus callosum spreading bihemispherically with dominance on the right side | ≈ 62 |
